# Knowledge-guided machine-learning and reverse screening combined method to predict cancer cell line responses to cytotoxic molecules

**DOI:** 10.64898/2025.12.16.694721

**Authors:** Alessandro Cuozzo, Marta A.S. Perez, Antoine Daina, Vincent Zoete

## Abstract

Estimating the cell line targets of cytotoxic small molecules is important for drug discovery and central for targeted therapy in oncology. Accurate prediction of sensitive cell lines enables early identification of efficacy and toxicity, optimization of drug selectivity, and can foster drug repurposing. While most bioactive compounds interact with multiple macromolecular targets, the cytotoxicity encompasses diverse complex biological and chemical mechanisms that could even not all be related to binding to macromolecules, making the prediction of cytotoxicity specificity particularly challenging. To address early-phase prediction of cancer cell line targets of cytotoxic compounds, we developed a method combining a machine-learning classification model with a ligand-based reverse screening procedure able to rank cell-line from the most probable to the least probable target of any cytotoxic molecule. The development focused on addressing the challenges related to the scarcity of available experimental data on non-cytotoxic compounds. A knowledge-guided generation of realistic alleged inactives allowed to train several binary logistic regression models. The most robust classification model was trained on 164,134 cytotoxic compounds extracted from ChEMBL to generate a score of predicted sensitivity of cell lines for any new cytotoxic molecule. The method demonstrated strong predictive ability, recovering at least one experimental target within the 15 most probable cell-lines for 71% of nearly 11,000 external cytotoxic compounds tested across 1018 cancer cell lines.

## 1. Introduction

The identification of cancer cell line targets of cytotoxic compounds represents a fundamental challenge in computer-aided drug design (CADD) in general and to support targeted therapy in oncology in particular^1^. Small molecules often interact with multiple macromolecules and cellular pathways, with effects varying dramatically across different cell lines due to genetic, epigenetic, and environmental heterogeneity. Early prediction of these cell line-specific interactions is critical for prioritizing compounds with desired therapeutic effects while identifying those likely to cause adverse effects, thereby reducing late-stage development failures and accelerating drug discovery^2,3^.

However, many cytotoxic compounds—whether classical chemotherapeutics or more recent targeted agents—still lack comprehensive characterization of their cell line specificity. Predictions of potential cancer cell line targets should therefore consider both categories of cytotoxic molecules: those with modes of actions making them ubiquitously cytotoxic and those having cell line-specific mechanisms.

Computational prediction of cancer cell line targets offers several practical advantages. It can steer experimental screening priorities, predict off-target effects in different cancer types, facilitate drug repurposing for accelerating drug development, and provide insights into compound selectivity toward different cell lines and classes.

While several methods have been successfully developed for predicting protein targets of bioactive molecules through ligand-based reverse screening^4,5^ , far fewer approaches specifically address cell line target prediction for cytotoxic compounds, and those that have been made available suffer from notable limitations.

Early efforts in cell line target prediction include CDRUG^6^, which estimated cytotoxicity probability using 8,565 active and 9,804 inactive compounds from the NCI-60 dataset, though the web tool is no longer available. Speck-Planche and colleagues developed many individual multi-target QSAR models to predict cytotoxicity against cancer cells from various tissues (prostate, breast, liver, brain, colorectal, connective tissue, urinary bladder), each trained on a few hundred molecules targeting 10 to 20 cell lines only^7–13^. However, these models aimed to predict if a molecule would be cytotoxic on all cell lines of a certain tissue without distinguishing between different cell lines from the same tissue or between cell lines annotated with the same disease. While potentially useful in the context of classical chemotherapies, such approaches are inadequate to support the design of targeted drugs.

ProTox-II^14^ predicts general cytotoxicity using random forests with diverse molecular fingerprints but was trained primarily on cytotoxic assays using the HepG2 liver cancer cell line as the only biological target. ChemBC^15^ built several hundreds of models using diverse molecular descriptors, fingerprints, and parameterizations, trained on 33,757 active and 21,152 inactive compounds against 13 breast cancer cell lines. Their best circular fingerprint-based random forest achieved accuracies of 0.84. Unfortunately, the website is not reachable anymore. Bonanni *et al*.^16^ developed ten machine-learning models focusing on two aggressive prostate cancer cell lines, achieving average accuracies in predicting compounds activity of 0.70 to 0.80 and Matthews Correlation Coefficients (MCC) of 0.50 to 0.60 for their most performing models.

Another tool, Cell-fishing^17^ achieved 66% accuracy on 758 cell lines but used a debatable fixed threshold of 10 μM to separate actives from inactives and relied on chemical structure similarity using a two-dimentsional description only.

CLC-Pred^18^ and its updated version CLC-Pred 2.0^19^ certainly represent the most comprehensive efforts in the field so far, predicting cytotoxicity against 391 cancer cell lines and 47 non-tumor cell lines using the PASS methodology^20^ . However, this method employs a data augmentation strategy, treating all compounds not tested against a given cell line as inactive—an approach that falsely classifies potentially ubiquitous agents like alkylating and intercalating agents as cell line-specific inactives, thus introducing substantial bias. Additionally, 70% of cell lines with prediction ability below 0.80 ROC AUC (area under the Receiver Operating Characteristic curve) were removed from the screenable set thus limiting the biological space.

Recently, deep learning attempts to incorporate multi-omics data, combining molecular descriptors with gene expression profiles, copy number variations, protein-protein interactions, and signaling pathways were published^21,22^. While powerful, these require additional experimental data not available for most compounds, hence suffering from restricted applicability domains. PaccMann^23^ developed by IBM for instance, achieved high performance in predicting IC_50_ of 135 kinase inhibitors on 1064 cell-line (R² = 0.86). The neural network being trained on less than 1,000 compounds, the method generalizability is limited, making its predictive capacity compromised.

The limitations of the methods developed so far developed are mainly due to the scarcity of highly curated experimental cytotoxicity data. In particular, a critical challenge in developing cell line target prediction models is the lack of experimentally confirmed inactive molecules. While databases like ChEMBL^24^ include extensive data on cytotoxic molecules genuinely non-cytotoxic compounds are comparatively very rare. This imbalance necessitates the generation of “alleged inactives“—compounds assumed inactive based primarily on absence of reported bioactivity. However, this strategy is especially problematic for cytotoxic agents for several reasons. Firstly, cytotoxic compounds can kill cells or inhibit their growth through different mechanisms, creating complexity that limits applicability of the similarity principle. This similarity principle states similar molecules are likely to exert comparable bioactivity is at the root of many predictive models in ligand-based drug design and quantified by us several time in the context of activity on protein targets^25,26^ . While similar cytotoxic molecules are expected to more likely target the same cell line, very different molecules can also target that cell line through multiple distinct mechanisms, potentially not all related to protein binding^27^. Therefore, randomly picking alleged inactives from the pool of known actives not tested on the biological target under investigation—a approached successfully employed for protein target prediction^5,20^ — becomes probably inappropriate for cell lines because different effective cytotoxic compounds tested and confirmed on different cell lines have significant probability to be active on other untested cell lines. Secondly, ubiquitous cytotoxic agents with mechanisms leading to cell-line non-specificity (*e.g.* DNA alkylation or acylation, DNA intercalation, antimetabolites, mitotic inhibition, and topoisomerase or gyrase inhibition) may be active on any cell line target, making it incorrect to assume they are inactive on untested cell lines.

We have developed a novel cell-line prediction method for cytotoxic compounds, inspired from our successful and thoroughly validated methodology for macromolecular target^5,28^, behind the SwissTargetPrediction webtool^29^. Accordingly, the presented approach aggregates a machine-learning classification model with a ligand-based virtual screening to generate a score of sensitivity enabling the ranking of the probable cell-line targets. The development focused on addressing the challenges dictated by the chemical and biological complexity of cytotoxicity and on the generation of relevant alleged inactives through refined knowledge-based strategies, while ensuring the model’s robustness, predictive ability and interpretability. The final model was trained on 164,134 compounds and achieved a success rate of 71% in correctly predicting at least one experimental target within the 15 most probable cell-lines for almost 11,000 external test compounds cytotoxic on 1018 cancer cell lines.

## 2. Methods

### 2.1 Data Collection and Curation

We extracted cytotoxicity data from ChEMBL^30^ version 29 using the following criteria: (i) small molecules with not more than 80 heavy atoms tested in cytotoxic or functional assays targeting cancer cell lines, (ii) cell lines unambiguously identified in Cellosaurus^31^ version 39 and not flagged as problematic or misidentified by ICLAC^32^ version 11, (iii) reported IC₅₀, EC₅₀, or GI₅₀ activity values, and (iv) human, mouse, rat, or dog cell lines. Each datapoint, defined as one compound-cell line interaction, was classified as active if the activity value is lower than 10 μM, inactive if the activity value exceeds 100 μM. Datapoints with activity values between 10 μM and 100 μM were considered as in a “grey area” and not used for the training.

The resulting *working dataset* contained 114,589 unique compounds tested experimentally against 1,048 cancer cell lines with at least two known actives (958 human, 68 mouse, 21 rat, 1 dog) from 51,777 assays, yielding 259,391 active datapoints, 28,209 inactive datapoints and 125,997 datapoints in the “grey area”. An *external test set* of 10,981 compounds was built by applying the same filtering criteria to ChEMBL version 32 and subsequently excluding compounds already within the working dataset.

### 2.2 Grouping Cell Lines by Lineage and Disease

Cell line information, in particular disease annotations, were retrieved and homogenized from Cellosaurus^31^, Cell Model Passports^33^, and the Cancer Cell Line Encyclopedia (CCLE)^34^. For CCLE, we considered primary disease, disease subtypes, and full disease classifications (primary disease + subtype). Whereas all cell lines had disease annotations in Cellosaurus, some lacked annotations in Cell Model Passports and/or CCLE. Missing annotations were inferred by generalizing Cellosaurus disease classifications according to corresponding annotations of other cell lines, with ambiguous cases manually curated by majority voting. Disease classifications were ultimately unified according to CCLE primary disease annotations, which classify cancers primarily by organ

### 2.3 Identification of potential and objective ubiquitous compounds

We developed chemical patterns encoded as SMARTS to identify *potential ubiquitous* compounds that may be cytotoxic across numerous cell lines because relying on cell line-unspecific mechanisms: Structural alerts from Begnini & Bossa^35^ as tabulated in OCHEM ToxAlerts^36^ and in SARpy^37^ were retrieved. This set of SMARTS was supplemented with patterns encoded by us, using MarvinSketch (version 22.7, www.chemaxon.com) and SMARTSviewer^38^ for molecules described in the medicinal chemistry literature as alkylating/acylating agents, intercalating agents, antimetabolites, mitotic inhibitors, and topoisomerase/gyrase inhibitors.^39–46^

All compiled patterns were tested against: (i) the set of bioactive compounds used to build SwissTargetPrediction (considered as negative control, since curated for *in vitro* activity on a well-defined protein targets)^29^ , (ii) the working set of the present study (section 2.1), and (iii) 169 FDA-approved or developmental non-specific anticancer drugs (54 alkylating/acylating agents, 17 intercalating agents, 56 antimetabolites, 22 mitotic inhibitors and 20 topoisomerase/gyrase inhibitors) gathered from DrugBank^47^, DrugCentral^48^, and Drugs.com (www.drugs.com) resources.

Moreover, we analyzed 10,567 PubMed abstracts (https://pubmed.ncbi.nlm.nih.gov/) referencing compounds in the working dataset (Section 2.1). In-house regular expressions were employed to verify that flagged categories of potential ubiquitous compounds (Supplementary Table 1) correspond to published mechanisms of cytotoxicity.

We filtered out SMARTS with regards to molecular structure of the compounds matching. Several patterns were excluded if it returned: i) more than 10,000 matches in SwissTargetPrediction set, ii) or less than 150 matches in working dataset (Section 2.1), or iii) no match in FDA-approved/developmental set. For redundant SMARTS, we privileged those with higher number of matches in ii) and iii), and then those that are more consistent with published mechanism (i.e. linked to more publications matching a regular expression that references the supposed category of the flagged compound). This process allowed to identify 13,731 compounds (12%) as potentially ubiquitous in our training set (Supplementary Table 2).

Finally, compounds experimentally active on more than 100 cell lines were considered as *objective ubiquitous* (79 compounds, Supplementary Figure 1).

### 2.4 Generation of alleged inactives

The limited number of genuinely inactive data points necessitated the generation of so-called *alleged* inactive data points.

Alleged inactives for each cell line were selected randomly from a pool of datapoints randomly until a ratio of 10 inactives per active was achieved (or approximated as closely as possible to this empirically defined ratio)^26^. A first pool of alleged inactive datapoints is thus produced assuming inactivity primarily because of the absence of reported bioactivity against a given biological target (here cancer cell lines, Figure 1A). Subsequent filters were applied to improve the definition of alleged inactive with further chemical knowledge (Figure 1B recognition of ubiquitous molecules, Figure 1C molecular scaffolds) and biological knowledge (Figure 1D disease associated to target cell-lines).

**Figure 1.**
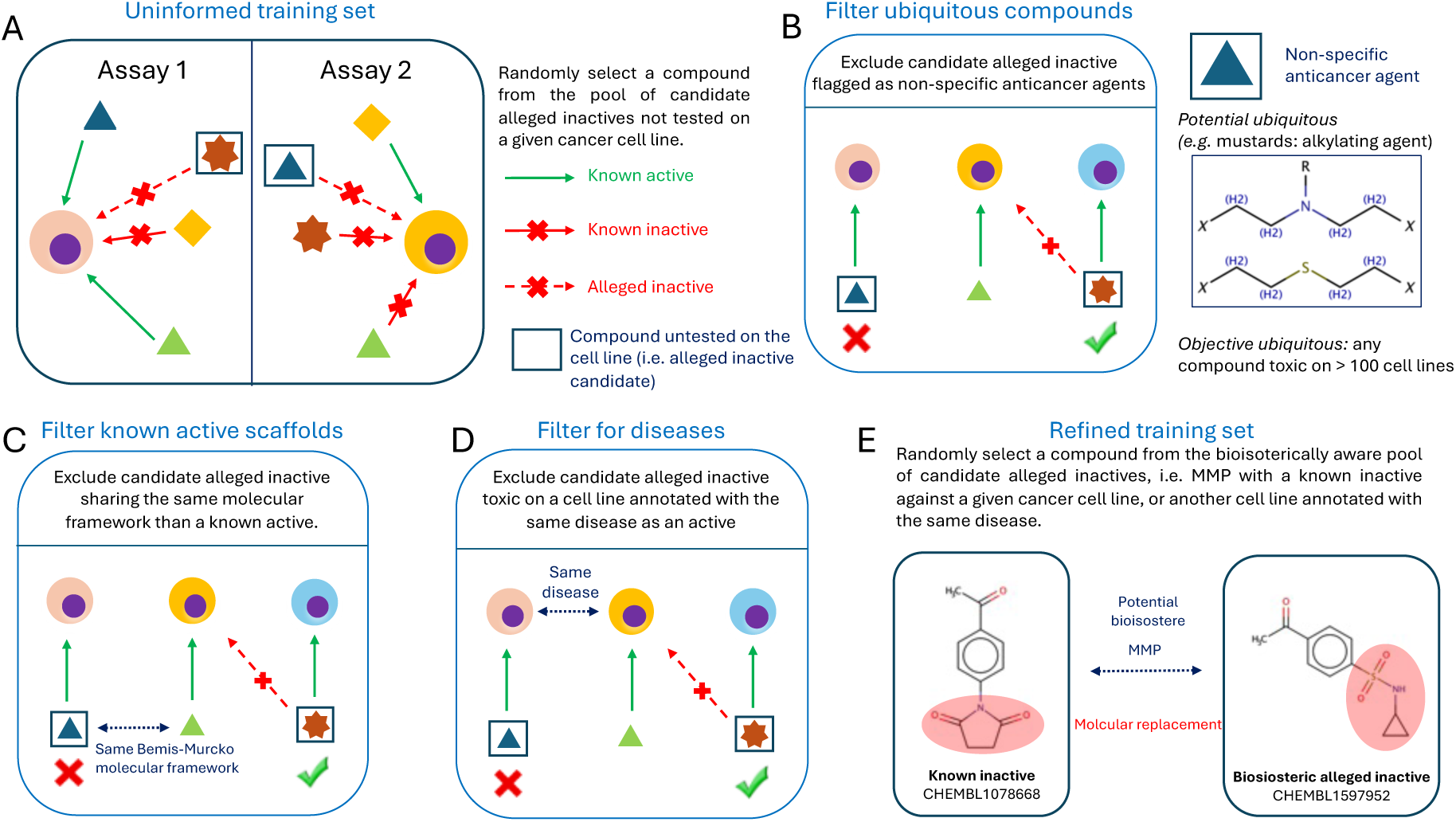
Strategies and filters to generate pools of alleged inactive datapoints. [A] The first *uninformed training set* was built with the pool of alleged inactives randomly selected from compounds untested experimentally among the cell line targets. Then knowledge-based filters were applied sequentially to refine the pool of alleged inactives by excluding compounds : [B] flagged as potential or objective ubiquitous cytotoxic molecules; [C] sharing the same Bemis-Murcko molecular framework with a known active on a given cell line; [D] experimentally cytotoxic on another cell line annotated with the same disease as the cell line target. [E] The final *refined training set* was built with another independent pool of alleged inactives randomly selected from compounds untested experimentally among the cell line targets and bioisosteres of these compounds, conducting a Matched Molecular Pair (MMP) analysis.

Another pool of alleged inactive datapoints was put together from possible bioisosteres of known inactives (Figure 1E) ; this collections of bioisosteric alleged inactive data point was also submitted to the three knowledge-based filters (Table 1).

**Table 1.**
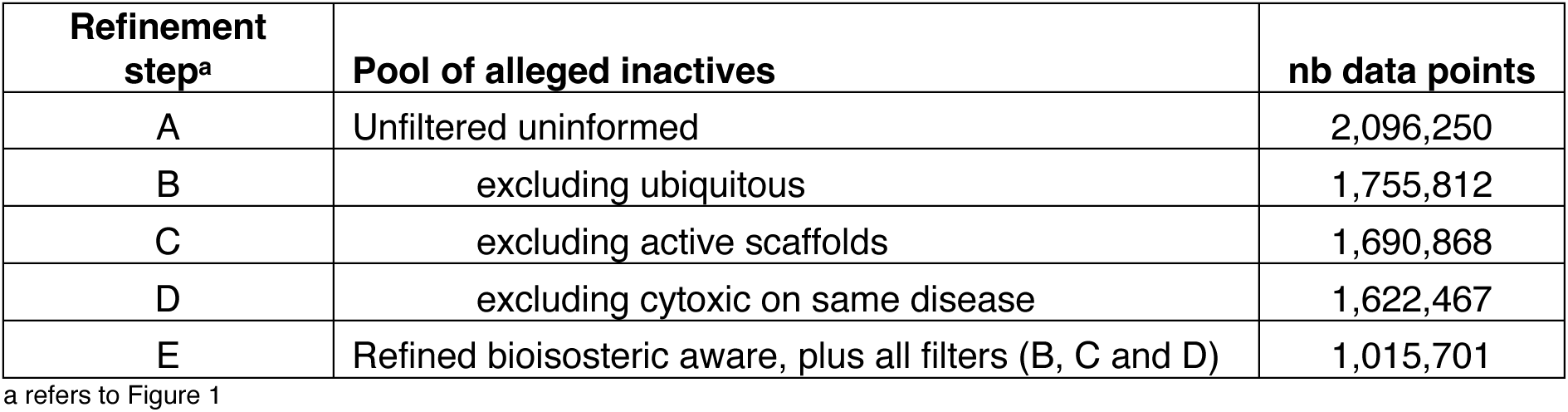
Generation of alleged inactive datapoint. pools by applying various knowledge-guided refinements.

Building on the SwissTargetPrediction methodology^5^ , alleged inactives for each cell line were randomly selected from compounds not tested on that cell line until a ratio of 10 inactives per active was achieved or approached (Figure 1A). 2,096,250 alleged inactive datapoints were thus generated to build a first dataset - involving 82’118 compounds experimentally assayed for cytotoxicity – and were added to 228’358 known active datapoints and 26’423 known inactive datapoints on 979 cancer cell lines. This uninformed training set contains 113’967 unique compounds in total.

In a second step (Figure 1B), compounds were excluded from the pool of alleged inactive candidates based on chemical knowledge, if they were previously flagged as (i) objective ubiquitous compounds (active on more than 100 cell lines) or (ii) potential ubiquitous compounds (flagged by chemical patterns, see section 2.3). At this step the training set contained 1,755,812 alleged inactive datapoints.

In a third step (Figure 1C), compounds were also excluded from the pool of alleged inactive candidates if sharing the same Bemis-Murcko molecular framework with any known active on the target cell line under consideration. After this filter, the pool of contained 1,690,868 alleged inactive datapoints.

The fourth step (Figure 1D) considered the disease annotations, by excluding from the above pool of alleged inactive candidate compounds if experimentally cytotoxic on at least one other cell line annotated with the same disease as the target cell line under consideration, according to CCLE primary disease classification. By applying this filter, 1,622,467 alleged inactive datapoints were in the pool.

Finally, we produced a new independent pool of alleged inactive candidates by selecting potential bioisosteric compounds of experimentally known inactives (Figure 1E). A matched molecular pair (MMP) analysis was conducted between all known inactives in the working dataset (as described in 2.1) and all ChEMBL 29 compounds untested in cytotoxicity assays. The MMP method is the one developed for the construction of the SwissBioisostere database.^49^ Here, we restricted the fragmentation to one single cut and the requirement that for being MMPs two compounds must differ by a molecular fragment with the same number of or less heavy atoms than the constant part of the molecules. 107,460 such bioisosteric compounds identified by MMP analysis were then further filtered and included in the pool of possible alleged inactives if they: (i) were not flagged as potential nor objective ubiquitous, (ii) did not share molecular frameworks with known actives, and (iii) had an MMP with a known inactive for the target cell line under consideration or for a cell line with the same disease annotation. The final number alleged inactive datapoints in the pool of candidates after applying all three filters described above reached 1,015,701. This final *refined training set* contained 164,134 compounds, involving 82,038 experimentally assayed for cytotoxicity on 836 cell lines and 82,096 as alleged inactives.

Both the *uninformed training set* and the *refined training set* were used independently to build binary logistic regression models for classification (see section 2.6). For the sake of analysis, intermediate filtered sets were also used to train the logistic regression.

### 2.5 Cheminformatics

Structural information for all compounds included in the working dataset and external test set as described in Sections 2.1 was stored in SMILES format^50^ after standardization (unsalting, desolvating, neutralizing, dearomatizing) with JChem Standardizer (version 21.3, www.chemaxon.com).

Every chemical structure was encoded in FP2 path-based fingerprints using OpenBabel^51^ (version 3.1.0, https://openbabel.org). Molecular frameworks were generated with RDKit (version 2023.03.2, www.rdkit.org) following the Bemis-Murcko definition^52^ .

The major tautomer and protonation state for each SMILES were obtained through the JChem Chemical calculations tool, then 20 conformations were produced by the JChem Structure manipulation tool (version 21.3, www.chemaxon.com) and stored in multi-MOL2 format.

For each conformation of each compound, the 3D-shape together with the projection of atomic electrostatic charges and of atomic contributions to lipophilicity was encoded as ES5D ElectroShape vectors^53,54^.

Similarity in chemical structure (2D_similarity_) of pairs of compounds was defined as the Tanimoto coefficient between FP2 bit strings. Shape similarity (3D_similarity_) was calculated according to (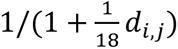), where *d_i,j_* is the smallest Manhattan distance between all 20 × 20 pairs of ES5D vectors.

### 2.6 Binary logistic model training and evaluation of robustness

Our methodology for training a binary logistic model for classification of probable biological target of small molecules has been extensively described elsewhere^5,26,29,55^.

In brief, two matrices were computed by pair-wise comparisons between all training compounds: one matrix including all 2D_similarity_ values and the other with all 3D_similarity_ values (see section 2.5).

Each training compound (active or inactive) was compared to all known actives of each cell line to retrieve the corresponding 2D_similarity_ and all 3D_similarity_ values. The highest values for each similarity metric become the features (2D_Score_ and 3D_Score_, respectively) to train the binary logistic regression for every cell line according to Equation 1.

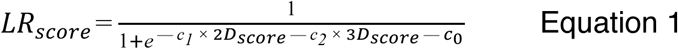

where *LR_score_* is a quantification of the probability of a given cell-line to be sensible to a cytotoxic compound (i.e. from training, “1” for known actives or “0” for known inactives on that cell line).

Models were evaluated through 10-fold cross-validation, with robustness of classification assessed using the Matthews Correlation Coefficient (MCC, Equation 2). True Positive Rate (TPR), True Negative Rate (TNR), Positive Predictive Value (PPV) and Negative Predictive Value (NPV) were also calculated (Equations 3, 4, 5, 6, respectively).

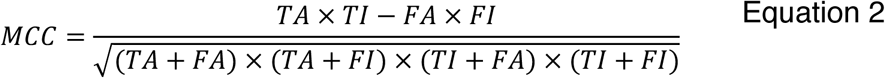

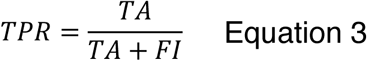

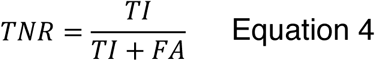

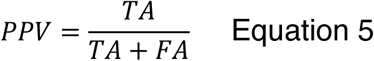

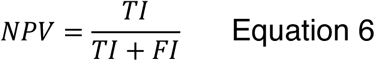

where, *TA* is the number of known actives returning *LR_score_* > 0.5; *TI* is the number of inactives returning *LR_score_* ≤ 0.5; *FA* is the number of inactives returning *LR_score_* > 0.5; *FI* is the number of known actives returning *LR_score_* ≤ 0.5

Training, cross-validation and MCC calculations were performed using the scikit-learn package^56^ (version 1.0.2, https://scikit-learn.org).

### 2.7 Membrane Permeation Prediction

Compound membrane permeation capacity of all compounds was estimated through a classification model to predict the parallel artificial membrane permeability assay (PAMPA) output developed by the National Center for Advancing Translational Sciences^57^. A probability of high permeation capacity is predicted for all compounds through a local installation of the PAMPA pH7.4 model, including Chemprop^58^ (version 1.1).

### 2.8 Reverse Screening and external validation

The reverse screening consisted in comparing any query molecule to all known actives on each screenable cell line to determine *2D_score_* and *3D_score_* values. The input of both features in the trained logistic equation led to an *LR_score* for each cell line. Cell lines are then ranked according *LR_score* values, from the most probably sensitive to the least probably sensitive to the query compound.

As external validation of our method, reverse screening was performed for the 10,981 compounds from the external test set (see Section 2.1). Success rates were calculated as the percentage of compounds for which at least one experimentally validated cell line target appeared in the top N predictions ranked according *LR_score* values (N = 1, 5, 15). Enrichment factors (EF) were computed as the ratio of success rate over random picking, where random picking is (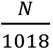), and 1,018 the number of screened cell lines.

Statistical significance of differences between compound categories (ubiquitous *vs*. non-ubiquitous, known *vs*. novel scaffolds, high *vs*. low permeability) was assessed using permutation tests with 1,000 random label shuffles, calculating P-values as the fraction of permutations leading to larger differences than observed on the unpermuted data set.

## 3. Results

### 3.1 Binary logistic model classification capacity

We assessed the influence on the classification capacity of our logistic model when applying successively each step of the strategy to generate alleged inactives and to build the training set (refer to Section 2.4). As shown in Figure 2A, the cross-validation MCC values increased progressively from the initial *uninformed training set* (0.61) through increasingly knowledge-informed approaches, reaching 0.83 for the model built on the final most *refined training*, including the selection of biosiosteric compounds to known inactives, along with the exclusion of potential and objective ubiquitous molecules, of molecules with scaffolds identical to known actives and of molecules with annotated activity on cell lines related to the same disease as known actives. Noteworthy, by following the SwissTargetPrediction methodology, training logistic sub-models after splitting the *refined training* set into 32 subsets based on query molecular sizes (ranging from less than 20 to more than 49 heavy atoms, Supplementary Figure 2) showed the same classification ability with an identical averaged cross-validated MCC of 0.83. This finding indicates that the model built on the entire *refined training* possesses sufficient robustness in its classification capability to encompass a range of drug-like small molecules, irrespective of their size.

**Figure 2.**
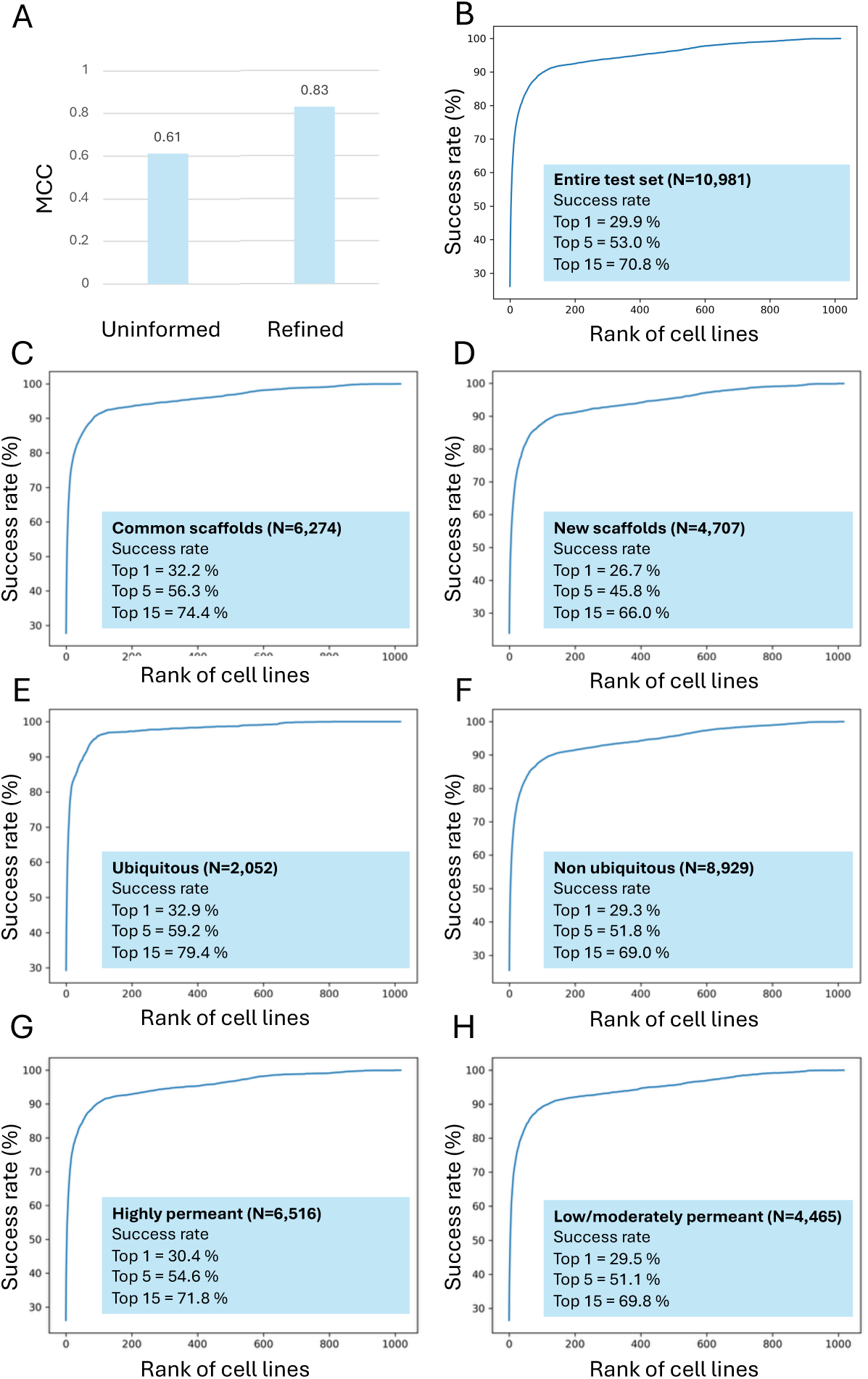
Robustness of the classification and predictive ability of the reverse screening. [A] 10-fold cross-validation Matthews Correlation Coefficient (MCC) of the logistic models for the first *Uninformed* to the final *Refined* training sets; Success rates in retrieving correctly one experimentally validated cell line target on [B] 10,981 cytotoxic test compounds external to training; [C] 6,274 test compounds bearing a common Bemis-Murcko molecular framework with training compounds; [D] 4,707 test compounds bearing a new Bemis-Murcko molecular framework (i.e. absent from training data); [E] 2,052 test compounds flagged as non-specific anticancer agents (i.e. potential ubiquitous compounds); [F] 8,929 test compounds with no potential ubiquitous flags. [G] 6,516 test compounds predicted with high permeation capacity; [H] 4,465 test compounds predicted with low or moderate permeation capacity.

The general improvement of classification ability principally resulted from increased TPR (See Supplementary Figure 3A), indicating better identification of actual experimentally validated cell line targets. The inclusion of negative bioisosteres, the exclusion of similar scaffolds to actives and of ubiquitous molecules appeared to have the most significant positive impact on the MCC, whereas the exclusion of compounds based on disease annotation has only a marginal effect on classification (See Supplementary Figure 3B)

### 3.2 External Validation on Novel Compounds

The predictive ability of our reverse screening method was evaluated on 10,981 compounds from ChEMBL version 32, external to the training set. The reverse screening coupled with the final logistic model built on the *refined training set* achieved 30% success rate on retrieving at least one experimentally validated cell line target at top 1, 53% at top 5, and 71% at top 15 (Figure 2B). This represents enrichment factors of 305, 108 and 48 at top1, top 5 and top respectively, compared to random picking.

A superior performance was found on the 6,274 test compounds sharing Bemis-Murcko with training data showed superior performance (32% at top 1, 56% at top 5, 74% at top 15; Figure 2C) compared to completely novel molecular frameworks (4707 compounds, 27% at top 1, 49% at top 5, 66% at top 15; Figure 2D). This relatively limited decrease of predictive ability for novel scaffolds (P < 0.005, permutation test) demonstrates the genuine capacity of the approach to reach new chemotypes, which is important for the generalization of the predictive method but also for scaffold-hopping—an efficient strategy to discover new chemical families towards specific biological targets (here, cell lines)^59^.

Higher success rates were obtained on the 2,052 test compounds flagged as potentially ubiquitous (33% at top 1, 59% at top 5, 79% at top 15; Figure 2E) than on the 8,929 non-ubiquitous compounds (29%, 52%, 69%; Figure 2F; P < 0.05). This likely reflects the higher probability that ubiquitous molecules are active on numerous cell lines (tested and untested), making experimental validated targets easier to find at the highest-ranked predicted cell lines. This finding reinforces the importance of identifying and appropriately handling non-specific ubiquitous anticancer agents for an unbiased interpretation of the reverse-screening output.

A trend can be observed about test compounds predicted as highly permeant showing slightly better performance (6,516 compounds Figure 2G) than those predicted with low/moderate permeability (4,465 compounds, Figure 2H), though differences were not statistically significant (P = 0.24 obtained through permutation test, Supplementary Figure 4C-E). This suggests that membrane permeation, which is much related to the mechanism of cytotoxicity and to the sub-cellular location of macromolecular targets, plays a role but is not the dominant factor in cell line selectivity.

### 3.3 Identification of Non-Specific Agents

The analysis of 10,567 PubMed abstracts confirmed correlation between our chemical pattern flags and published mechanism descriptions, validating our identification strategy and the criteria for selecting the relevant patterns (as described in Section 2.3.

Our chemical pattern collection identified 17,449 potential ubiquitous compounds in the working dataset (15.2%) and 6,403 in the test data set (19.2%). The majority of which (84.6%) were flagged as alkylating, acylating, or intercalating agents.

Interestingly, potential bioisosteres of known inactives in the final refined training set showed similar frequency of ubiquitous compounds (16.9%), but with fewer intercalating agents and more antimetabolites in proportion. This is consistent with bioisosteres being derived from discovery compounds across the whole ChEMBL database, most of which not being specifically designed for cytotoxicity.

### 3.5 Cell Line Grouping by Disease

Grouping cell lines by CCLE primary disease annotation proved effective, enabling alleged inactive generation for cell lines that would otherwise have insufficient data. This high-quality annotation balances specificity (distinguishing various cancer types) with tractability (enough cell lines per group), categorizing diseases primarily by organ while treating exceptionally aggressive tumors separately (refer to (Supplementary Table 3 for details). The 32 primary disease groups showed heterogeneous data distribution, with lung cancer having the most cell lines (185) and known actives (22,712), while some rare cancers had single cell lines (Supplementary Figure 5).

## 4. Discussion

The efficiency of the described method, combining a machine-learning logistic regression model with ligand-based reverse screening, is demonstrated by its strong predictive capacity on an external test set. Specifically, for 30% of the 10,981 cytotoxic compounds not included in the training set, the method correctly identified at least one experimentally validated cell line as the most probable target (Figure 2B). This performance represents a remarkable 305-fold enrichment compared to random picking among the 1018 screenable cell lines. The success rate for the top 5 and top 15 predicted cell lines reached 53% and 71%, respectively.

Overall, our predictive method demonstrates substantial improvements over previously published approaches. For example, CLC-Pred 2.0^19^ reported a mean accuracy of prediction of 0.9 across diverse datasets and setups (defined as ROC AUC) but its performance relied on a questionable data augmentation by treating all untested compounds as inactive. In contrast, our approach is more conservative by applying stricter data curation and our knowledge-based refined strategy to generate realistic alleged inactives prove to markedly ameliorates the classification ability of our logistic regression model. Moreover, 70% of cell lines with poor model performance were removed from today’s available version of CLC-pred. With 1,018 predictable cell-line across 32 cancer types, our screening set covers a much larger space of biological targets. This holds patently true when comparing with ChemBC^15^, whose best random-forest classification models on circular fingerprints achieved high accuracy (0.84) but on 13 breast cancer cells, only. Another tool named Cell-fishing^17^ achieved 66% accuracy on 758 cell lines but used a questionable 10 μM activity cutoff and relied solely on 2D similarity. By contrast, our cytotoxicity data curation sets more rigorous thresholds (<10 μM cytotoxic, >100 μM non-cytotoxic, buffered by a “gray area” in between) providing better discrimination between known actives and inactives. Also, our cell-line sensitivity prediction relies on a machine-learning method combining similarity metrics in 2D and in 3D, it allows a better generalization with a larger applicability domain and the findings of different chemotypes^26^. Furthermore, our reverse-screening methodology outputs a quantitative score (LR_score_). While this predictive value is not the strict numerical probability of the sensitivity of a cell-line, it enables the ranking of more than 1000 cancer cell-lines as possible targets for a given cytotoxic compound.

The predictive ability of the presented method is not at the same level as similar approaches applied to protein target prediction. As an example, the reverse-screening behind SwissTargetPrediction reached success rates at 51% at top 1, 65% at top and 73% at top 15 in similar validation exercises, but on larger test sets^26^. One cause is certainly the more limited amount of experimental cytotoxicity data compared to *in vitro* bioactivity on well-defined proteins. However as well, this indicates the peculiarities of the modes of action of cytotoxicity that make the biological processes less tightly related to the similarity principle. Several of these factors such as putative multiple protein targets, non-specific mode of cytotoxicity possibly unrelated to macromolecule binding, access to the subcellular location of macromolecular target(s) were considered in the development of our methods, especially in the knowledge-based generation of alleged inactives, whose amount and quality are crucial to balance the scarcity of available experimentally validated non-cytotoxic molecules and to train an effective predictive model.

Our bioisosterically aware strategy for producing alleged inactives, based on MMP, tackles two major challenges in cell line target prediction: the scarcity of experimentally confirmed inactive compounds and the complexity of cytotoxic mechanisms. A classical simple random selection of alleged inactives by inferring that a molecule not reported as active on a given biological target can probably be an inactive, as successfully employed for protein target prediction, is problematic for cell lines. This is because compounds active compounds on one cell line have more chances to be active on many others, a phenomenon particularly true for ubiquitous cytotoxic agents.

Besides, we found that generating potential bioisosteres of known inactives have two additional benefits. Firstly, the membrane permeability capacity (or non-capacity) is conserved because bioisosteric replacements typically maintain similar physicochemical properties, including those governing membrane permeation capacity (like lipophilicity or polarity)^60^ . This is supported by the observation that the alleged inactives generated through the MMP analysis are in average less permeant than the alleged inactives obtained by traditional random selection (Table 2, 47.0% and 56.4% of compounds predicted with high permeation capacity, respectively). Moreover, the 47.0% of predicted permeant compounds obtained within the MMP-derived molecules is close to the 43.0% of known inactives. In contrast, the 56.4% of permeant traditional alleged inactive is closer to the property of cytotoxic compounds globally (54.8% to 59.4%) or even to any bioactive molecules (SwissTargetPrediction and ChEMBL 29 data 53.3% and 53.6%, respectively). Secondly, while keeping the physicochemical profile constant, the bioisosteric analysis identifies compounds from different chemotypes (novel Bemis-Murcko scaffolds). This reduces overfitting to specific chemical series and improves generalization within an extended chemical space.

**Table 2.** Prediction of membrane permeation capacity for different types of compounds in the dataset.

|  | # compounds | high predicted permeation capacity |
| --- | --- | --- |
| Working dataset<sup>a</sup> | 114,546 | 54.8 % |
| Known actives | 77,853 | 55.9 % |
| Known inactives | 4,600 | 43.0 % |
| Traditional alleged inactives<sup>b</sup> | 83,704 | 56.4 % |
| External test set | 10,981 | 59.4 % |
| SwissTargetPrediction | 482,094 | 53.3 % |
| ChEMBL 29<sup>c</sup> | 1,917,558 | 53.5 % |
| Bioisosteric alleged inactives<sup>d</sup> | 107,460 | 47.0 % |

a see section 2.1
b randomly picked among the known actives
c compounds from ChEMBL 29, excluding those contained in the main dataset
d see section 2.4

The statistically significant higher success rate for ubiquitous versus non-ubiquitous compounds (Figure 2E-F; P < 0.05) highlights a key aspect of non-specific cytotoxic agents’ behavior that must be considered separately from cell-selective cytotoxic molecules. Ubiquitous compounds (being alkylating/acylating agents, intercalating agents, antimetabolites, mitotic inhibitors or topoisomerase/gyrase inhibitors) displayed higher success not because they are better predicted, but because they are genuinely active on numerous cell lines—including probably many untested ones. Flagging such ubiquitously cytotoxic molecules is important for proper interpretation of the output of our method. Indeed, the success rates for non-specific agents reflects more a statistical bias, due to the inability to follow the similarity principle, rather than pure predictive accuracy. In practice, if a compound is detected by our pattern recognition process, it is recommended to disregard the quantification of probability, and instead to consider a quantitative assessment. Indeed, this detection is useful to support drug discovery since ubiquitous molecule may have broader spectrum of cytotoxicity leading to a more acute risk of unwanted effects and should be handled differently in drug research pipelines (if not directly removed in most cases).

The envisaged practical applications of this novel predictive method echo its screening-oriented nature. Indeed, high enrichment helps to narrow substantially experimental screening requirements. This paves the way to efficient prioritization of cell lines for experimental phenotypic oncology screening. The ability to predict targets for novel scaffolds (e.g. with 70% success rate at top 15) enables scaffold hopping to discover novel cytotoxic chemotypes. In addition, thanks to the curated disease annotations, identifying new cell line targets for existing drugs can foster development for unmet medical needs by drug repurposing.

However, several limitations of our new predictive method should be considered. Our collection of chemical patterns to detect cytotoxic ubiquitous compounds, while comprehensive, was optimized for current data and may require expansion as novel chemotypes or novel drug classes emerge.

While PAMPA prediction was considered, the expected relationship with cytotoxicity was weaker than anticipated. This may reflect the complexity of cellular uptake mechanisms or limitations in current permeability models. Future work will evaluate refined predictions considering not only passive permeation but active transport and subcellular localization of potential protein targets. The presented method is able to predict cell line targets but not mechanisms of cytotoxicity. The perspective of crossing its output with prediction of protein targets (*e.g.* using SwissTargetPrediction) is expected to provide useful mechanistic insights.

While predicting off-target toxicity on cancer cell lines provides insights into compound selectivity, including bioactivity data from other non-cancer cell lines could be useful to help beholding global safety and toxicity of the molecules under study. Besides, integerating information on genomic alterations in specific cell lines with compound sensitivity predictions could support treatment selection for personalized oncology. Finally, the same methodology could be applied to estimate cytotoxicity on other species. In particular, application to microorganisms could be of relevance for the discovery of novel antibacterial, antimycotic or antiviral treatments. The absolute requisite will be the availability of data of sufficient quality and quantity for a model and a screening set to be built.

## 4. Conclusion

We developed a machine learning model based on binary logistic regression able to classify possible cancer cell line targets of cytotoxic compounds by combining 2D and 3D molecular similarity metrics. The knowledge-driven strategy to generate negative training data points (*alleged inactives*) and the meticulous experimental data curation allowed efficient training and led to robust classification for 164,134 cytotoxic compounds (MCC = 0.83 in 10-fold cross-validation). The sensitivity score obtained enabled the ranking of 1018 screenable cancer cell lines from the most to the least probable target of any cytotoxic small molecule.

The predictive ability of the whole method was demonstrated by the success rate of the reverse virtual screening of an external test set of 10,981 cytotoxic compounds. One experimental cell-line target was correctly predicted within the 15 predicted most probable in 70% of the cases. The 48-fold enrichment observed is indicative of the practicability of the approach in supporting cell-line prioritization in experimental phenotypic screening in the field of oncology. In such experimental settings, the deconvolution of macromolecular targets and the deciphering of modes of action are central points. The feasibility to address these problematics will be assessed by future studies crossing the protein target predictions obtained by SwissTargetPrediction and the output of the here presented methods.

## 5. Acknowledgement

The authors would like to thank Professor Elisa Oricchio at EPFL, for the crucial discussions on the biology of cancer cell-lines. Chemaxon (www.chemaxon.com) is acknowledged for the licensing agreement.

## Notes

### Competing Interest Statement

The authors have declared no competing interest.

## Bibliographic references

1. Prasse, P., et al. Matching anticancer compounds and tumor cell lines by neural networks with ranking loss. NAR Genom. Bioinform. 4, lqab128 (2022).

2. Basith, S., Cui, M., Macalino, S. J. Y. & Choi, S. Expediting the Design, Discovery and Development of Anticancer Drugs using Computational Approaches. Curr. Med. Chem. 24, 1–26 (2018).

3. Cui, W. et al. Discovering Anti-Cancer Drugs via Computational Methods. Front. Pharmacol. 11, 733 (2020).

4. Keiser, M. J. et al. Relating protein pharmacology by ligand chemistry. Nat. Biotechnol. 25, 197–206 (2007).

5. Gfeller, D., Michielin, O. & Zoete, V. Shaping the interaction landscape of bioactive molecules. Bioinformatics 29, 3073–3079 (2013).

6. Li, G.-H. & Huang, J.-F. CDRUG: a web server for predicting anticancer activity of chemical compounds. Bioinformatics 28, 3334–3335 (2012).

7. Speck-Planche, A., Kleandrova, V. V., Luan, F. & Cordeiro, M. N. D. S. Multi-target drug discovery in anti-cancer therapy: Fragment-based approach toward the design of potent and versatile anti-prostate cancer agents. Bioorg. Med. Chem. 19, 6239–6244 (2011).

8. Speck-Planche, A., Kleandrova, V. V., Luan, F. & Cordeiro, M. N. D. S. Chemoinformatics in anti-cancer chemotherapy: Multi-target QSAR model for the in silico discovery of anti-breast cancer agents. European Journal of Pharmaceutical Sciences 47, 273–279 (2012).

9. Kleandrova, V. V., Scotti, M. T., Scotti, L., Nayarisseri, A. & Speck-Planche, A. Cell-based multi-target QSAR model for design of virtual versatile inhibitors of liver cancer cell lines. SAR QSAR Environ. Res. 31, 815–836 (2020).

10. Speck-Planche, A., Kleandrova, V. V., Luan, F. & Cordeiro, M. N. D. S. Chemoinformatics in multi-target drug discovery for anti-cancer therapy: in silico design of potent and versatile anti-brain tumor agents. Anti-cancer agents Med. Chem. 12, 678–85 (2011).

11. Speck-Planche, A., Kleandrova, V. V., Luan, F. & Cordeiro, M. N. D. S. Rational drug design for anti-cancer chemotherapy: Multi-target QSAR models for the in silico discovery of anti-colorectal cancer agents. Bioorg. Med. Chem. 20, 4848–4855 (2012).

12. Speck-Planche, A., Kleandrova, V. V., Luan, F. & Cordeiro, M. N. D. S. Fragment-based QSAR model toward the selection of versatile anti-sarcoma leads. Eur. J. Med. Chem. 46, 5910–5916 (2011).

13. Speck-Planche, A., Kleandrova, V. V., Luan, F. & Cordeiro, M. N. D. S. Unified multi-target approach for the rational in silico design of anti-bladder cancer agents. Anti-cancer agents Med. Chem. 13, 791–800 (2012).

14. Banerjee, P., Eckert, A. O., Schrey, A. K. & Preissner, R. ProTox-II: a webserver for the prediction of toxicity of chemicals. Nucleic Acids Res. 46, W257–W263 (2018).

15. He, S. et al. Machine Learning Enables Accurate and Rapid Prediction of Active Molecules Against Breast Cancer Cells. Front. Pharmacol. 12, 796534 (2021).

16. Bonanni, D., Pinzi, L. & Rastelli, G. Development of machine learning classifiers to predict compound activity on prostate cancer cell lines. J. Cheminformatics 14, 77 (2022).

17. Tejera, E. et al. Cell fishing: A similarity based approach and machine learning strategy for multiple cell lines-compound sensitivity prediction. PLoS ONE 14, e0223276 (2019).

18. Lagunin, A. A. et al. CLC-Pred: A freely available web-service for in silico prediction of human cell line cytotoxicity for drug-like compounds. PLoS ONE 13, e0191838–13 (2018).

19. Lagunin, A. A. et al. CLC-Pred 2.0: A Freely Available Web Application for In Silico Prediction of Human Cell Line Cytotoxicity and Molecular Mechanisms of Action for Druglike Compounds. Int. J. Mol. Sci. 24, 1689 (2023).

20. Poroikov, V., Filimonov, D., Lagunin, A., Gloriozova, T. & Zakharov, A. PASS: identification of probable targets and mechanisms of toxicity†. SAR QSAR Environ Res 18, 101–110 (2007).

21. Baptista, D., Ferreira, P. G. & Rocha, M. Deep learning for drug response prediction in cancer. Brief. Bioinform. 22, 360–379 (2020).

22. Zhang, H., Chen, Y. & Li, F. Predicting Anticancer Drug Response With Deep Learning Constrained by Signaling Pathways. Front. Bioinform. 1, 639349 (2021).

23. Cadow, J., Born, J., Manica, M., Oskooei, A. & Martínez, M. R. PaccMann: a web service for interpretable anticancer compound sensitivity prediction. Nucleic Acids Res. 48, W502–W508 (2020).

24. Mendez, D. et al. ChEMBL: towards direct deposition of bioassay data. Nucleic Acids Res. 47, D930–D940 (2019).

25. Bragina, M. E., Daina, A., Perez, M. A. S., Michielin, O. & Zoete, V. The SwissSimilarity 2021 Web Tool: Novel Chemical Libraries and Additional Methods for an Enhanced Ligand-Based Virtual Screening Experience. Int J Mol Sci 23, 811 (2022).

26. Daina, A. & Zoete, V. Testing the predictive power of reverse screening to infer drug targets, with the help of machine learning. Commun. Chem. 7, 105 (2024).

27. Galluzzi, L. et al. Molecular mechanisms of cell death: recommendations of the Nomenclature Committee on Cell Death 2018. Cell Death Differ. 25, 486–541 (2018).

28. Basant, N., Gupta, S. & Singh, K. P. Predicting binding affinities of diverse pharmaceutical chemicals to human serum plasma proteins using QSPR modelling approaches. SAR QSAR Environ Res 27, 67–85 (2016).

29. Daina, A., Michielin, O. & Zoete, V. SwissTargetPrediction: updated data and new features for efficient prediction of protein targets of small molecules. Nucleic Acids Res. 47, W357–W364 (2019).

30. Zdrazil, B. et al. The ChEMBL Database in 2023: a drug discovery platform spanning multiple bioactivity data types and time periods. Nucleic Acids Res. 52, D1180–D1192 (2023).

31. Bairoch, A. The Cellosaurus, a Cell-Line Knowledge Resource. J Biomol Tech jbt.18-2902-002 (2018) doi:10.7171/jbt.18-2902-002.

32. Capes-Davis, A. et al. Check your cultures! A list of cross-contaminated or misidentified cell lines. Int. J. Cancer 127, 1–8 (2010).

33. Meer, D. van der et al. Cell Model Passports-a hub for clinical, genetic and functional datasets of preclinical cancer models. Nucleic Acids Res. 47, D923–D929 (2019).

34. Nusinow, D. P. et al. Quantitative Proteomics of the Cancer Cell Line Encyclopedia. Cell 180, 387–402.e16 (2020).

35. Benigni, R. & Bossa, C. Structural Alerts of Mutagens and Carcinogens. Curr. Comput. Aided-Drug Des. 2, 169–176 (2006).

36. Sushko, I., et al. Online chemical modeling environment (OCHEM): web platform for data storage, model development and publishing of chemical information. J Comput Aided Mol Des 25, 533–554 (2011).

37. Ferrari, T. et al. Automatic knowledge extraction from chemical structures: the case of mutagenicity prediction. SAR QSAR Environ. Res. 24, 365–383 (2013).

38. Ehrt, C., Krause, B., Schmidt, R., Ehmki, E. S. R. & Rarey, M. SMARTS.plus - A Toolbox for Chemical Pattern Design. Mol. Inform. 39, e2000216 (2020).

39. Froelich-Ammon, S. J. & Osheroff, N. Topoisomerase Poisons: Harnessing the Dark Side of Enzyme Mechanism (∗). J. Biol. Chem. 270, 21429–21432 (1995).

40. Kluge, A. F. & Petter, R. C. Acylating drugs: redesigning natural covalent inhibitors. Curr. Opin. Chem. Biol. 14, 421–427 (2010).

41. Lansiaux, A. Les antimétabolites. Bull. du Cancer 98, 1263–1274 (2011).

42. Luengo, A., Gui, D. Y. & Heiden, M. G. V. Targeting Metabolism for Cancer Therapy. Cell Chem. Biol. 24, 1161–1180 (2017).

43. Martino, E. et al. Vinca alkaloids and analogues as anti-cancer agents: Looking back, peering ahead. Bioorg. Med. Chem. Lett. 28, 2816–2826 (2018).

44. Mosca, L., Ilari, A., Fazi, F., Assaraf, Y. G. & Colotti, G. Taxanes in cancer treatment: Activity, chemoresistance and its overcoming. Drug Resist. Updat. 54, 100742 (2021).

45. Pommier, Y. Topoisomerase I inhibitors: camptothecins and beyond. Nat. Rev. Cancer 6, 789–802 (2006).

46. Venugopal, S., Sharma, V., Mehra, A., Singh, I. & Singh, G. DNA intercalators as anticancer agents. Chem. Biol. Drug Des. 100, 580–598 (2022).

47. Wishart, D. S. et al. DrugBank 5.0: a major update to the DrugBank database for 2018. Nucleic Acids Res. 46, D1074–D1082 (2018).

48. Halip, L. et al. Exploring DrugCentral: from molecular structures to clinical effects. J. Comput.-Aided Mol. Des. 37, 681–694 (2023).

49. Cuozzo, A., Daina, A., Perez, M. A. S., Michielin, O. & Zoete, V. SwissBioisostere 2021: updated structural, bioactivityand physicochemical data delivered by a reshapedweb interface. Nucleic Acids Res 50, D1382–D1390 (2021).

50. Weininger, D. SMILES, a chemical language and information system. 1. Introduction to methodology and encoding rules. J. Chem. Inf. Comput. Sci. 28, 31–36 (1988).

51. O’Boyle, N. M. et al. OpenBabel: An open chemical toolbox. J. Cheminform. 3, 33 (2011).

52. Bemis, G. W. & Murcko, M. A. The properties of known drugs. 1. Molecular frameworks. J. Med. Chem. 39, 2887–2893 (1996).

53. Armstrong, M. S. et al. ElectroShape: fast molecular similarity calculations incorporating shape, chirality and electrostatics. J Comput Aided Mol Des 24, 789–801 (2010).

54. Armstrong, M. S., Finn, P. W., Morris, G. M. & Richards, W. G. Improving the accuracy of ultrafast ligand-based screening: incorporating lipophilicity into ElectroShape as an extra dimension. J Comput Aided Mol Des 25, 785–790 (2011).

55. Gfeller, D. et al. SwissTargetPrediction: a web server for target prediction of bioactive small molecules. Nucleic Acids Res. 42, W32–W38 (2014).

56. Pedregosa, F. et al. Scikit-learn: Machine Learning in Python. Journal of Machine Learning Research 12, 2825–2830 (2011).

57. Siramshetty, V. et al. Validating ADME QSAR Models Using Marketed Drugs. SLAS Discov. 26, 1326–1336 (2021).

58. Heid, E. et al. Chemprop: a machine learning package for chemical property prediction. Journal of Chemical Information and Modeling 64, 9–17 (2023).

59. Schneider, G., Neidhart, W., Giller, T. & Schmid, G. “Scaffold-Hopping” by Topological Pharmacophore Search: A Contribution to Virtual Screening. Angew. Chem. Int. Ed. Engl. 38, 2894–2896 (1999).

60. Daina, A. & Zoete, V. A BOILED-Egg To Predict Gastrointestinal Absorption and Brain Penetration of Small Molecules. ChemMedChem 11, 1117–1121 (2016).

