## Supplementary material for "Knowledge-guided machine-learning and reverse screening combined method to predict cancer cell line responses to cytotoxic molecules"

Supplementary Table 1 | In-house regular expressions associated with non-specific anticancer agent category

Abstracts of scientific publications referencing cytotoxic compounds retrieved from experimental assays were analysed. Compounds were labeled as potentially belonging to a category of non-specific anticancer agent if the abstract contained one or several associated regular expressions (right column, underlying rows).

<table>
<tr><td>acylating agents</td><td></td></tr>
<tr><td></td><td>(acylat|alkanoylat)</td></tr>
<tr><td></td><td></td></tr>
<tr><td>alkylating agents</td><td></td></tr>
<tr><td></td><td>(alkylat|ethylat)</td></tr>
<tr><td></td><td>(the){0,1} {0,1}(double){0,1}[ -]{0,1}(strand){0,1}(ed){0,1}[ -<br>]{0,1}(helical){0,1}(helix){0,1}(helicoid(al){0,1}){0,1}(genomic){0,1}(core){0,1}(mitochondrial){0,1}(<br>nucleus|nucleic){0,1}s{0,1}'{0,1} {0,1}(dna|base[ -]pair|(acid){0,1}[ -<br>]{0,1}deoxyribonucleic|nucleotide|nucleic acid|m(in|aj)or<br>groove|gene|exon|intron|exome|introme|genome|purine|pyrimidine)[ -](alkyl|ehtyl|methyl)</td></tr>
<tr><td></td><td>(the){0,1} {0,1}(double){0,1}[ -]{0,1}(strand){0,1}(ed){0,1}[ -<br>]{0,1}(helical){0,1}(helix){0,1}(helicoid(al){0,1}){0,1}(genomic){0,1}(core){0,1}(mitochondrial){0,1}(<br>nucleus|nucleic){0,1}s{0,1}'{0,1} {0,1}(dna|base[ -]pair|(acid){0,1}[ -<br>]{0,1}deoxyribonucleic|nucleotide|nucleic acid|m(in|aj)or<br>groove|gene|exon|intron|exome|introme|genome|purine|pyrimidine)[ -<br>](attack|crush|dismantle|impair|gut|consume|eradicate|destroy|annihilate|<br>end|terminate|smash|break|damage|bind|link|stick|insert|interpolate|inject|intersperse|int<br>roduce|add|adduct|interject|interpose|edge|insinuate|work|cut|fit|weave|sandwich|inset|<br>append|install|inlay|interfile|wedge|interline|thrust|attach|lard|shove|cram)(ed|s|ing){0,1}<br>{0,1}(in|to|into|for){0,1} {0,1}(the){0,1}</td></tr>
<tr><td></td><td>(attack|crush|dismantle|impair|gut|consume|eradicate|destroy|annihilate|<br>end|terminate|smash|break|damage|bind|link|stick|insert|interpolate|inject|intersperse|int<br>roduce|add|adduct|interject|interpose|edge|insinuate|work|cut|fit|weave|sandwich|inset|<br>append|install|inlay|interfile|wedge|interline|thrust|attach|lard|shove|cram)(ed|s|ing){0,1}<br>{0,1}(in|to|into|for){0,1} {0,1}(the){0,1}[ -<br>]{0,1}(themselves|itself|afar|aside|alongside){0,1}(between|between<br>in|betwixt|amid|amidst|among|at intervals of|halfway|in|to|into|in the middle|in the<br>midst|in the seam|in the thick|medially|mid|midway|within){0,1}( of){0,1}[ -]{0,1}(the){0,1}<br>{0,1}(cancer|cancerous|tumor){0,1}[ -]{0,1}(cell){0,1}(ular){0,1}(s){0,1}[ -]{0,1}(line){0,1}s{0,1}<br>{0,1}(type){0,1}s{0,1} {0,1}s{0,1}'{0,1}[ -]{0,1}(the){0,1} {0,1}(double){0,1}[ -<br>]{0,1}(strand){0,1}(ed){0,1}[ -<br>]{0,1}(helical){0,1}(helix){0,1}(helicoid(al){0,1}){0,1}(genomic){0,1}(core){0,1}(mitochondrial){0,1}(<br>nucleus|nucleic){0,1}s{0,1}'{0,1} {0,1}(dna|base[ -]pair|(acid){0,1}[ -<br>]{0,1}deoxyribonucleic|nucleotide|nucleic acid|m(in|aj)or<br>groove|gene|exon|intron|exome|introme|genome|purine|pyrimidine)</td></tr>
<tr><td></td><td>(between|between in|betwixt|amid|amidst|among|at intervals of|halfway|in|to|into|in the<br>middle|in the midst|in the seam|in the thick|medially|mid|midway|within)( of){0,1}[ -<br>]{0,1}(the){0,1} {0,1}(cancer|cancerous|tumor){0,1}[ -]{0,1}(cell){0,1}(ular){0,1}(s){0,1}[ -<br>]{0,1}(line){0,1}s{0,1} {0,1}(type){0,1}s{0,1} {0,1}s{0,1}'{0,1}(the){0,1} {0,1}(double){0,1}[ -<br>]{0,1}(strand){0,1}(ed){0,1}[ -<br>]{0,1}(helical){0,1}(helix){0,1}(helicoid(al){0,1}){0,1}(genomic){0,1}(core){0,1}(mitochondrial){0,1}(<br>nucleus|nucleic){0,1}s{0,1}'{0,1} {0,1}(dna|base[ -]pair|(acid){0,1}[ -<br>]{0,1}deoxyribonucleic|nucleotide|nucleic acid|m(in|aj)or<br>groove|gene|exon|intron|exome|introme|genome|purine|pyrimidine)</td></tr>
<tr><td></td><td></td></tr>
<tr><td>intercalating agents</td><td></td></tr>
<tr><td></td><td>(anti|inhibit|stop|halt|impede|hinder|hamper|obstruct|embarrass|stymie|handicap|restrain<br>|block|delay|disrupt|constrain|encumber|cramp|thwart|tie<br>up|hobble|fetter|clog|shackle|short[ -]{0,1}circuit|handcuff|trammel|stifle|hold<br>back|frustrate|curb|hog[ -<br>]{0,1}tie|bind|rein|interfere|cramp|manacle|retard|retain|tie|brake|give a hard<br>time|balk|chain|hold<br>up|strangle|confine|sabotage|arrest|tether|derail|check|barricade|blockade|repress|roadbl<br>ock|baffle|halter|leash|choke|foil|suppress|smother|suffocate|muzzle|stump|mire|bog|hed<br>ge|hem) {0,1}(ors{0,1} of|with|down|in|ing|for){0,1} {0,1}(the){0,1} {0,1}(proper|correct){0,1}</td></tr>
</table>

<table>
<tr><td></td><td>{0,1}(good|normal|standard|usual){0,1} {0,1}(use|usage|utility|function(ing){0,1}){0,1} {0,1}(of){0,1} {0,1}(topoisomerase|gyrase)</td></tr>
<tr><td></td><td>(topoisomerase|gyrase)[ -](anti|inhibit|stop|halt|impede|hinder|hamper|obstruct|embarrass|stymie|handicap|restrain|block|delay|disrupt|constrain|encumber|cramp|thwart|tie up|hobble|fetter|clog|shackle|short[ -]{0,1}circuit|handcuff|trammel|stifle|hold back|frustrate|curb|hog[ -]{0,1}tie|bind|rein|interfere|cramp|manacle|retard|retain|tie|brake|give a hard time|balk|chain|hold up|strangle|confine|sabotage|arrest|tether|derail|check|barricade|blockade|repress|roadblock|baffle|halter|leash|choke|foil|suppress|smother|suffocate|muzzle|stump|mire|bog|hedge|hem) {0,1}(ors{0,1} of|with|down|in|ing|for){0,1} {0,1}(the){0,1} {0,1}(proper|correct){0,1} {0,1}(good|normal|standard|usual){0,1} {0,1}(use|usage|utility|function(ing){0,1}){0,1} {0,1}(of){0,1}</td></tr>
<tr><td></td><td>intercalat</td></tr>
<tr><td></td><td>(the){0,1} {0,1}(double){0,1}[ -]{0,1}(strand){0,1}(ed){0,1}[ -]{0,1}(helical){0,1}(helix){0,1}(helicoid(al){0,1}){0,1}(genomic){0,1}(core){0,1}(mitochondrial){0,1}(nucleus|nucleic){0,1}s{0,1}'{0,1} {0,1}(dna|base[ -]pair|(acid){0,1}[ -]{0,1}deoxyribonucleic|nucleotide|nucleic acid|m(in|aj)or groove|gene|exon|intron|exome|introme|genome|purine|pyrimidine)[ -](attack|crush|dismantle|impair|gut|consume|eradicate|destroy|annihilate|end|terminate|smash|break|damage|bind|link|stick|insert|interpolate|inject|intersperse|introduce|add|adduct|interject|interpose|edge|insinuate|work|cut|fit|weave|sandwich|inset|append|install|inlay|interfile|wedge|interline|thrust|attach|lard|shove|cram)(ed|s|ing){0,1} {0,1}(in|to|into|for){0,1} {0,1}(the){0,1}</td></tr>
<tr><td></td><td>(attack|crush|dismantle|impair|gut|consume|eradicate|destroy|annihilate|end|terminate|smash|break|damage|bind|link|stick|insert|interpolate|inject|intersperse|introduce|add|adduct|interject|interpose|edge|insinuate|work|cut|fit|weave|sandwich|inset|append|install|inlay|interfile|wedge|interline|thrust|attach|lard|shove|cram)(ed|s|ing){0,1} {0,1}(in|to|into|for){0,1} {0,1}(the){0,1}[ -]{0,1}(themselves|itself|afar|aside|alongside){0,1}(between|between in|betwixt|amid|amidst|among|at intervals of|halfway|in|to|into|in the middle|in the midst|in the seam|in the thick|medially|mid|midway|within){0,1}( of){0,1}[ -]{0,1}(the){0,1} {0,1}(cancer|cancerous|tumor){0,1}[ -]{0,1}(cell){0,1}(ular){0,1}(s){0,1}[ -]{0,1}(line){0,1}s{0,1} {0,1}(type){0,1}s{0,1} {0,1}s{0,1}'{0,1}[ -]{0,1}(the){0,1} {0,1}(double){0,1}[ -]{0,1}(strand){0,1}(ed){0,1}[ -]{0,1}(helical){0,1}(helix){0,1}(helicoid(al){0,1}){0,1}(genomic){0,1}(core){0,1}(mitochondrial){0,1}(nucleus|nucleic){0,1}s{0,1}'{0,1} {0,1}(dna|base[ -]pair|(acid){0,1}[ -]{0,1}deoxyribonucleic|nucleotide|nucleic acid|m(in|aj)or groove|gene|exon|intron|exome|introme|genome|purine|pyrimidine)</td></tr>
<tr><td></td><td>(between|between in|betwixt|amid|amidst|among|at intervals of|halfway|in|to|into|in the middle|in the midst|in the seam|in the thick|medially|mid|midway|within)( of){0,1}(the){0,1} {0,1}(double){0,1}[ -]{0,1}(strand){0,1}(ed){0,1}[ -]{0,1}(helical){0,1}(helix){0,1}(helicoid(al){0,1}){0,1}(genomic){0,1}(core){0,1}(mitochondrial){0,1}(nucleus|nucleic){0,1}s{0,1}'{0,1} {0,1}(dna|base[ -]pair|(acid){0,1}[ -]{0,1}deoxyribonucleic|nucleotide|nucleic acid|m(in|aj)or groove|gene|exon|intron|exome|introme|genome|purine|pyrimidine)</td></tr>
<tr><td></td><td></td></tr>
<tr><td>antimetabolites</td><td></td></tr>
<tr><td></td><td>(target|attack|aim|anti|inhibit|stop|halt|impede|hinder|hamper|obstruct|embarrass|stymie|handicap|restrain|block|delay|disrupt|constrain|encumber|cramp|thwart|tie up|hobble|fetter|clog|shackle|short[ -]{0,1}circuit|handcuff|trammel|stifle|hold back|frustrate|curb|hog[ -]{0,1}tie|bind|rein|interfere|cramp|manacle|retard|retain|tie|brake|give a hard time|balk|chain|hold up|strangle|confine|sabotage|arrest|tether|derail|check|barricade|blockade|repress|roadblock|baffle|halter|leash|choke|foil|suppress|smother|suffocate|muzzle|stump|mire|bog|hedge|hem) {0,1}(ors{0,1} of|with|down|in|ing|for){0,1} {0,1}(the){0,1} {0,1}(proper|correct){0,1} {0,1}(good|normal|standard|usual){0,1} {0,1}(use|usage|utility|function(ing){0,1}){0,1} {0,1}(of){0,1}[ -]{0,1}(the){0,1} {0,1}(cancer|cancerous|tumor){0,1}[ -]{0,1}(cell){0,1}(ular){0,1}(s){0,1}[ -]{0,1}(line){0,1}s{0,1} {0,1}(type){0,1}s{0,1} {0,1}s{0,1}'{0,1}[ -]{0,1}metaboli</td></tr>
<tr><td></td><td>metaboli(c|te(s){0,1}|sm)[ -](target|attack|aim|anti|inhibit|stop|halt|impede|hinder|hamper|obstruct|embarrass|stymi</td></tr>
</table>

<table>
<tr><td></td><td>e|handicap|restrain|block|delay|disrupt|constrain|encumber|cramp|thwart|tie up|hobble|fetter|clog|shackle|short[ -]{0,1}circuit|handcuff|trammel|stifle|hold back|frustrate|curb|hog[ -]{0,1}tie|bind|rein|interfere|cramp|manacle|retard|retain|tie|brake|give a hard time|balk|chain|hold up|strangle|confine|sabotage|arrest|tether|derail|check|barricade|blockade|repress|roadblock|baffle|halter|leash|choke|foil|suppress|smother|suffocate|muzzle|stump|mire|bog|hedge|hem) {0,1}(ors{0,1} of|with|down|in|ing|for){0,1} {0,1}(the){0,1} {0,1}(proper|correct){0,1} {0,1}(good|normal|standard|usual){0,1} {0,1}(use|usage|utility|function(ing){0,1}){0,1} {0,1}(of){0,1}</td></tr>
<tr><td></td><td></td></tr>
<tr><td>mitotic inhibitors</td><td></td></tr>
<tr><td></td><td>(microtubule|tubulin|(cyto|cell(ular){0,1}[ -]{0,1}skelet))</td></tr>
<tr><td></td><td>(target|attack|aim|anti|inhibit|stop|halt|impede|hinder|hamper|obstruct|embarrass|stymie|handicap|restrain|block|delay|disrupt|constrain|encumber|cramp|thwart|tie up|hobble|fetter|clog|shackle|short[ -]{0,1}circuit|handcuff|trammel|stifle|hold back|frustrate|curb|hog[ -]{0,1}tie|bind|rein|interfere|cramp|manacle|retard|retain|tie|brake|give a hard time|balk|chain|hold up|strangle|confine|sabotage|arrest|tether|derail|check|barricade|blockade|repress|roadblock|baffle|halter|leash|choke|foil|suppress|smother|suffocate|muzzle|stump|mire|bog|hedge|hem) {0,1}(ors{0,1} of|with|down|in|ing|for){0,1} {0,1}(the){0,1} {0,1}(proper|correct){0,1} {0,1}(good|normal|standard|usual){0,1} {0,1}(use|usage|utility|function(ing){0,1}){0,1} {0,1}(of){0,1} {0,1}(the){0,1} {0,1}(cancer|cancerous|tumor){0,1}[ -]{0,1}(cell){0,1}(ular){0,1}(s){0,1}[ -]{0,1}(line){0,1}s{0,1} {0,1}(type){0,1}s{0,1} {0,1}s{0,1}'{0,1} {0,1}(mitosis|mitotic|cycle|division|divid|separat|segregat)</td></tr>
<tr><td></td><td>(add to|appreciate|augment|boost|build up|complement|enlarge|heighten|increase|intensify|raise|reinforce|strengthen|upgrade|adorn|aggrandize|amplify|beautify|boom|elevate|embroider|exaggerate|exalt|lift|magnify|pad|pyramid|swell|flesh out|enhance|provoke|trigger|induce|force|cause|prompt|set off|spark|set in motion|activate|give rise to) {0,1}(rs{0,1} of){0,1}(s|ed|ing){0,1} {0,1}(of){0,1} {0,1}(the){0,1} {0,1}(the){0,1} {0,1}(cancer|cancerous|tumor){0,1}[ -]{0,1}(cell){0,1}(ular){0,1}(s){0,1}[ -]{0,1}(line){0,1}s{0,1} {0,1}(type){0,1}s{0,1} {0,1}s{0,1}'{0,1} senescence</td></tr>
<tr><td></td><td>(the){0,1} {0,1}(cancer|cancerous|tumor){0,1}[ -]{0,1}(cell){0,1}(ular){0,1}(s){0,1}[ -]{0,1}(line){0,1}s{0,1} {0,1}(type){0,1}s{0,1} {0,1}s{0,1}'{0,1} {0,1}(mitosis|mitotic|cycle|division|divid|separat|segregat)[ -](target|attack|aim|anti|inhibit|stop|halt|impede|hinder|hamper|obstruct|embarrass|stymie|handicap|restrain|block|delay|disrupt|constrain|encumber|cramp|thwart|tie up|hobble|fetter|clog|shackle|short[ -]{0,1}circuit|handcuff|trammel|stifle|hold back|frustrate|curb|hog[ -]{0,1}tie|bind|rein|interfere|cramp|manacle|retard|retain|tie|brake|give a hard time|balk|chain|hold up|strangle|confine|sabotage|arrest|tether|derail|check|barricade|blockade|repress|roadblock|baffle|halter|leash|choke|foil|suppress|smother|suffocate|muzzle|stump|mire|bog|hedge|hem) {0,1}(ors{0,1} of|with|down|in|ing|for){0,1} {0,1}(the){0,1} {0,1}(proper|correct){0,1} {0,1}(good|normal|standard|usual){0,1} {0,1}(use|usage|utility|function(ing){0,1}){0,1} {0,1}(of){0,1}</td></tr>
<tr><td></td><td>(the){0,1} {0,1}(cancer|cancerous|tumor){0,1}[ -]{0,1}(cell){0,1}(ular){0,1}(s){0,1}[ -]{0,1}(line){0,1}s{0,1} {0,1}(type){0,1}s{0,1} {0,1}s{0,1}'{0,1} senescence[ -](add to|appreciate|augment|boost|build up|complement|enlarge|heighten|increase|intensify|raise|reinforce|strengthen|upgrade|adorn|aggrandize|amplify|beautify|boom|elevate|embroider|exaggerate|exalt|lift|magnify|pad|pyramid|swell|flesh out|enhance|provoke|trigger|induce|force|cause|prompt|set off|spark|set in motion|activate|give rise to) {0,1}(rs{0,1} of){0,1}(s|ed|ing){0,1} {0,1}(of){0,1} {0,1}(the){0,1}</td></tr>
<tr><td></td><td></td></tr>
<tr><td>topoisosmerase/gyrase inhibitors</td><td></td></tr>
<tr><td></td><td>(topoisomerase|gyrase)</td></tr>
</table>

Supplementary Table 2 | SMARTS chemical used to detect potential ubiquitous cytotoxic compounds

Categories of non-specific anticancer agents (left column) contain one or several ubiquitous flags (middle column, underlying rows). Ubiquitous flags are assigned to a query molecule if the latter contains one or multiple related SMARTS patterns (right column, underlying rows).

| alkyating agents | | |
|---|---|---|
| | S_or_N_mustard_TA362 | |
| | | [F,Cl,Br,I][CH2][CH2][NX3,SX2][CH2][CH2][F,Cl,Br,I] |
| | S_or_N_mustard_SA5 | |
| | | CCNCCCl |
| | Aliphatic_halogens_TA365 | |
| | | [CX4;!H0][Br,Cl,I] |
| | Epoxides_and_aziridines_TA364 | |
| | | [CX4]1[OX2,NX3][CX4]1 |
| | Quinones_TA369 | |
| | | [$([#6X3]1=,:[#6X3]-,:[#6X3](=[OX1])-,:[#6X3]=,:[#6X3]-,:[#6X3]1(=[OX1])),$([#6X3]1(=[OX1])-,:[#6X3](=[OX1])-,:[#6X3]=,:[#6X3]-,:[#6X3]=,:[#6X3]1)] |
| | Nitrosoureas_TA790 | |
| | | [NX3]C(=O)[$([NH](C=O)[NX2]=O),$(N(C=O)([#6])[NX2]=O)] |
| | Oxazaphosphorine | |
| | | [$([#1,*])]-[#7](-[$([#1,*])])P1(=O)[#8]-[#6]-[#6]-[#6]-[#7]1-[$([#1,*])] |
| | azide_or_triazene_group_SA22 | |
| | | NN=N |
| | | N=[N+]=[N-] |
| | Azide_and_triazene_groups_TA379 | |
| | | [$([NX2!R]=[NX2!R][NX3!R]),$([NX2]=[NX2+]=[NX1-]),$([NX2]=[NX2+]=N)] |
| | Hydrazine_TA370 | |
| | | [NX3;!$([NX3](=[OX1])=[OX1]);!$([NX3+](=[OX1])[O-])][NX3;!$([NX3](=[OX1])=[OX1]);!$([NX3+](=[OX1])[O-])] |
| | Alkyl_and_aryl_N-nitroso_groups_TA378 | |
| | | [#6][NX3][NX2]=[OX1] |
| | alkyl_(C<5)_or_benzyl_ester_of_sulfonic_or_phosphonic_acid_SA2 | |
| | | S(=O)(=O)(OCC) |

| | | |
|---|---|---|
| | Alkyl_or_benzyl_ester_of_sulphonic_or_phosphonic_acid_TA359 | |
| | | [$([Sv6X4;!$([Sv6X4][OH]);!$([Sv6X4][SH]);!$([Sv6X4][O-]);!$([Sv6X4][S-])](=[OX1])(=[OX1])[$([OX2]C([#1,F,Cl,Br,I])([#1,F,Cl,Br,I])[#1,F,Cl,Br,I]),$([OX2]C([#1,F,Cl,Br,I])([#1,F,Cl,Br,I])C([#1,F,Cl,Br,I])([#1,F,Cl,Br,I])[#1,F,Cl,Br,I]),$([OX2]C([#1,F,Cl,Br,I])([#1,F,Cl,Br,I])C([#1,F,Cl,Br,I])([#1,F,Cl,Br,I])C([#1,F,Cl,Br,I])([#1,F,Cl,Br,I])[#1,F,Cl,Br,I]),$([OX2]C([#1,F,Cl,Br,I])(C([#1,F,Cl,Br,I])([#1,F,Cl,Br,I])[#1,F,Cl,Br,I])C([#1,F,Cl,Br,I])([#1,F,Cl,Br,I])[#1,F,Cl,Br,I]),$([OX2]C([#1,F,Cl,Br,I])([#1,F,Cl,Br,I])C([#1,F,Cl,Br,I])([#1,F,Cl,Br,I])C([#1,F,Cl,Br,I])([#1,F,Cl,Br,I])C([#1,F,Cl,Br,I])([#1,F,Cl,Br,I])[#1,F,Cl,Br,I]),$([OX2]C([#1,F,Cl,Br,I])(C([#1,F,Cl,Br,I])([#1,F,Cl,Br,I])[#1,F,Cl,Br,I])C([#1,F,Cl,Br,I])([#1,F,Cl,Br,I])C([#1,F,Cl,Br,I])([#1,F,Cl,Br,I])[#1,F,Cl,Br,I]),$([OX2][CH0](C([#1,F,Cl,Br,I])([#1,F,Cl,Br,I])[#1,F,Cl,Br,I])(C([#1,F,Cl,Br,I])([#1,F,Cl,Br,I])[#1,F,Cl,Br,I])C([#1,F,Cl,Br,I])([#1,F,Cl,Br,I])[#1,F,Cl,Br,I]),$([OX2]C([#1,F,Cl,Br,I])([#1,F,Cl,Br,I])C([#1,F,Cl,Br,I])(C([#1,F,Cl,Br,I])([#1,F,Cl,Br,I])[#1,F,Cl,Br,I])C([#1,F,Cl,Br,I])([#1,F,Cl,Br,I])[#1,F,Cl,Br,I]),$([OX2][CH2]c1ccccc1)]),$([Pv5X4;!$([Pv5X4][OH]);!$([Pv5X4][SH]);!$([Pv5X4][O-]);!$([Pv5X4][S-])](=[OX1])([$([OX2]C([#1,F,Cl,Br,I])([#1,F,Cl,Br,I])[#1,F,Cl,Br,I]),$([OX2]C([#1,F,Cl,Br,I])([#1,F,Cl,Br,I])C([#1,F,Cl,Br,I])([#1,F,Cl,Br,I])[#1,F,Cl,Br,I]),$([OX2]C([#1,F,Cl,Br,I])([#1,F,Cl,Br,I])C([#1,F,Cl,Br,I])([#1,F,Cl,Br,I])C([#1,F,Cl,Br,I])([#1,F,Cl,Br,I])[#1,F,Cl,Br,I]),$([OX2]C([#1,F,Cl,Br,I])(C([#1,F,Cl,Br,I])([#1,F,Cl,Br,I])[#1,F,Cl,Br,I])C([#1,F,Cl,Br,I])([#1,F,Cl,Br,I])[#1,F,Cl,Br,I]),$([OX2]C([#1,F,Cl,Br,I])([#1,F,Cl,Br,I])C([#1,F,Cl,Br,I])([#1,F,Cl,Br,I])C([#1,F,Cl,Br,I])([#1,F,Cl,Br,I])C([#1,F,Cl,Br,I])([#1,F,Cl,Br,I])[#1,F,Cl,Br,I]),$([OX2]C([#1,F,Cl,Br,I])(C([#1,F,Cl,Br,I])([#1,F,Cl,Br,I])[#1,F,Cl,Br,I])C([#1,F,Cl,Br,I])([#1,F,Cl,Br,I])C([#1,F,Cl,Br,I])([#1,F,Cl,Br,I])[#1,F,Cl,Br,I]),$([OX2][CH0](C([#1,F,Cl,Br,I])([#1,F,Cl,Br,I])[#1,F,Cl,Br,I])(C([#1,F,Cl,Br,I])([#1,F,Cl,Br,I])[#1,F,Cl,Br,I])C([#1,F,Cl,Br,I])([#1,F,Cl,Br,I])[#1,F,Cl,Br,I]),$([OX2]C([#1,F,Cl,Br,I])([#1,F,Cl,Br,I])C([#1,F,Cl,Br,I])(C([#1,F,Cl,Br,I])([#1,F,Cl,Br,I])[#1,F,Cl,Br,I])C([#1,F,Cl,Br,I])([#1,F,Cl,Br,I])[#1,F,Cl,Br,I]),$([OX2][CH2]c1ccccc1)])[$([OX2]C([#1,F,Cl,Br,I])([#1,F,Cl,Br,I])[#1,F,Cl,Br,I]),$([OX2]C([#1,F,Cl,Br,I])([#1,F,Cl,Br,I])C([#1,F,Cl,Br,I])([#1,F,Cl,Br,I])[#1,F,Cl,Br,I]),$([OX2]C([#1,F,Cl,Br,I])([#1,F,Cl,Br,I])C([#1,F,Cl,Br,I])([#1,F,Cl,Br,I])C([#1,F,Cl,Br,I])([#1,F,Cl,Br,I])[#1,F,Cl,Br,I]),$([OX2]C([#1,F,Cl,Br,I])(C([#1,F,Cl,Br,I])([#1,F,Cl,Br,I])[#1,F,Cl,Br,I])C([#1,F,Cl,Br,I])([#1,F,Cl,Br,I])[#1,F,Cl,Br,I]),$([OX2]C([#1,F,Cl,Br,I])([#1,F,Cl,Br,I])C([#1,F,Cl,Br,I])([#1,F,Cl,Br,I])C([#1,F,Cl,Br,I])([#1,F,Cl,Br,I])C([#1,F,Cl,Br,I])([#1,F,Cl,Br,I])[#1,F,Cl,Br,I]),$([OX2]C([#1,F,Cl,Br,I])(C([#1,F,Cl,Br,I])([#1,F,Cl,Br,I])[#1,F,Cl,Br,I])C([#1,F,Cl,Br,I])([#1,F,Cl,Br,I])C([#1,F,Cl,Br,I])([#1,F,Cl,Br,I])[#1,F,Cl,Br,I]),$([OX2][CH0](C([#1,F,Cl,Br,I])([#1,F,Cl,Br,I])[#1,F,Cl,Br,I])(C([#1,F,Cl,Br,I])([#1,F,Cl,Br,I])[#1,F,Cl,Br,I])C([#1,F,Cl,Br,I])([#1,F,Cl,Br,I])[#1,F,Cl,Br,I]),$([OX2]C([#1,F,Cl,Br,I])([#1,F,Cl,Br,I])C([#1,F,Cl,Br,I])(C([#1,F | 

| | | |
|---|---|---|
| | | ,Cl,Br,I])([#1,F,Cl,Br,I])[#1,F,Cl,Br,I])C([#1,F,Cl,Br,I])([#1,F,Cl,Br,I])[#1,F,Cl,Br,I]),$([OX2][CH2]c1ccccc1)])] |
| | Simple_aldehyde_TA368 | |
| | | [CX3]([H])(=[OX1])[#1,#6&!$([CX3]=[CX3])] |
| | Alkyl_sulfonate | |
| | | [#6]-[#8]S([#6])(=O)=O |
| | Ethylene_imine | |
| | | [$([#1,*])]-[#6]-1-[#6](-[$([#1,*])])-[#7]-1-[$([#1,*])] |
| | alpha_beta_unsaturated_aliphatic_alkoxy_group_SA24 | |
| | | C=[CH]O |
| | Monohaloalkene_TA361 | |
| | | [CX3]([CX4,#1])([F,Cl,Br,I])=[CX3]([CX4,#1])[!F!Cl!Br!I] |
| | α_β-Unsaturated_carbonyl_TA367 | |
| | | [CX3]([!$([OH]);!$([O-])])](=[OX1])[CX3H1]=[CX3]([$([CH3]),$([CH2][CH3]),$([CH2][CH2][CH3]),$([CH]([CH3])[CH3]),$([CH2][CH2][CH2][CH3]),$([CH]([CH3])[CH2][CH3]),$([CH2][CH]([CH3])[CH3]),$([CH0]([CH3])([CH3])[CH3]),$([CH2][CH2][CH2][CH2][CH3]),$([CH]([CH3])[CH2][CH2][CH3]),$([CH2][CH]([CH3])[CH2][CH3]),$([CH2][CH2][CH]([CH3])[CH3]),$([CH]([CH2][CH3])[CH2][CH3]),$([CH]([CH3])[CH]([CH3])[CH3]),$([CH0]([CH3])([CH3])[CH2][CH3]),$([CH2][CH0]([CH3])([CH3])[CH3]),$([#1,#7,#8,F,Cl,Br,I,#15,#16,#5]),$([CH]=[CH][#6]);!$([a!r0])])[$([CH3]),$([CH2][CH3]),$([CH2][CH2][CH3]),$([CH]([CH3])[CH3]),$([CH2][CH2][CH2][CH3]),$([CH]([CH3])[CH2][CH3]),$([CH2][CH]([CH3])[CH3]),$([CH0]([CH3])([CH3])[CH3]),$([CH2][CH2][CH2][CH2][CH3]),$([CH]([CH3])[CH2][CH2][CH3]),$([CH2][CH]([CH3])[CH2][CH3]),$([CH2][CH2][CH]([CH3])[CH3]),$([CH]([CH2][CH3])[CH2][CH3]),$([CH]([CH3])[CH]([CH3])[CH3]),$([CH0]([CH3])([CH3])[CH2][CH3]),$([CH2][CH0]([CH3])([CH3])[CH3]),$([#1,#7,#8,F,Cl,Br,I,#15,#16,#5]),$([CH]=[CH][#6]);!$([a!r0])] |
| | Duocarmycin_derivative | |
| | | [#6]@[#6]@[#7](@[#6]-,=,:1-,=,:[#6]-,=,:[#6]-,=,:[#6]-,=,:[#6]-,=,:[#6]-,=,:1)-[#6](=O)-[#6]1-,=,:[#6]c2ccccc2[#7]1 |
| | Distamycin_or_netropsin_derivative | |
| | | [$([#1,*])]-[#6]-[#7](-[$([#1,*])])-[#6](=O)-[#6]1=,:[#6][#6](=,:[#6][#7]1-[$([#1,*])])-[#7](-[$([#1,*])])-[#6](-[$([#1,*])])=O |
| | Platinum | |
| | | [Pt]([$([#1,*])])[$([#1,*])] |

| | | |
|---|---|---|
| intercalating agents | | |
| | 3_or_more_fused_aromatic_rings | |
| | | a:1:a:a:2:a:a:a:3:a:a:a:a:a:3:a:2:a:1 |
| | | a:1:a:2:a:a:a:a:a:2:a:2:a:a:a:a:a:1:2 |
| | | a:1:a:a:2:a:a:a:a:3:a:a:a:a(:a:1):a:2:3 |
| | | a:1:a:a:a:2:a:a:3:a:a:a:a:a:3:a:a:2:a:1 |
| | | a:1:a:a:a:2:a(:a:1):a:a:a:1:a:a:a:a:a:2:1 |
| | | a:1:a:a:2:a:3:a:1:a:a:a:a:3:a:a:1:a:a:a:a:a:2:1 |
| | | a:1:a:a:a:2:a(:a:1):a:1:a:a:a:a:3:a:a:a:a:2:a:1:3 |
| | | a:1:a:a:2:a:a:a:3:a:a:a:a:4:a:a:a(:a:1):a:2:a:3:4 |
| | | a:1:a:2:a:a:a:a:a:2:a:2:a:1:a:a:a:1:a:a:a:a:a:2:1 |
| | | *-1-*:2:*:*:*:*:*:2-*:2:*:*:*:*:*-1:2 |
| | | *-1-*:2:*:*:*:*:*:2-*:2:*:*:3:*:*:*:*:*:3:*:*-1:2 |
| | | *-1-*:2:*:*:*:*:*:2-*:2:*-1:*:*:*:1:*:*:*:*:*:2:1 |
| | | *-1-*-*:2:*:*:*:*:3:*:*:*:*(-*-1):*:2:3 |
| | | *-1-*:2:*:*:*:*:*:2-*-*:2:*:*:*:*:*-1:2 |
| | | *:1:*:*:2:*:*:3:*:*:*:*:*:3:*:*:2:*:1 |
| | | *:1:*:*:2:*:*:3:*:*:*:*:3:*:*:2:*:1 |
| | | *:1:*:*:2:*:*:*:3:*:*:*:*:3:*:2:*:1 |
| | | *:1:*:*:2:*:*:*:*:3:*:*:*:*:1:*:2:3 |
| | Anthraquinone | |
| | | O=[#6]-1-c2ccccc2-[#6](=O)-c2ccccc-12 |
| | Piperidine-2,6-dione | |
| | | [$([#1,*])]-[#7]-1-[#6](=O)-c2cccc3cccc(-[#6]-1=O)c23 |
| acylating agents | | |
| | Isocyanate_and_isothiocyanate_groups_TA372 | |
| | | [NX2]=[CX2]=[OX1,Sv2X1] |
| antimetabolites | | |
| | Purine_analog | |
| | | [#7,#8,#16]-,=[#6]~1~[#7;X3H1,X2H0]~[#6](-[#1,#7,#9,#17])~[#7;X3H1,X2H0]~[#6]~2~[#6]~1~[#7;X3H1,X2H0]~[#6]~[#7]~2-[$([#1,*])] |
| | | [#8]-,=[#6]~1~[#6]~[#7;X3H1,X2H0]~[#6]~[#7;X3H1,X2H0]~[#6]~2~[#6]~1~[#7;X3H1,X2H0]~[#6]~[#7]~2-[$([#1,*])] |

| | | |
|---|---|---|
| | Pyrimidine_analog | |
| | | [#7,#8]-,=[#6]~1~[#7;X3H1,X2H0]~[#6](=O)~[#7](-[#1,*])~[#6;X4H2,X3H1]~[#6]~1-[#9,#6,#1] |
| | Antifolate | |
| | | [#7]-[#6]~1~[#7]~[#6]~[#6]~2~[#6](~[#6]~[#7]~[#6]~2~[#7]~1)~[#6]~1~[#6]~[#6]~[#6]~[#6,#7,#8,#16]~1 |
| | | [#7]-[#6]~1~[#6]~[#6]~[#6]~2~[#6]~[#6](~[#6]~[#6]~[#6]~2~[#6]~1)~[#6]~1~[#6]~[#6]~[#6]~[#6,#7,#8,#16]~1 |
| | | [#7]-[#6]~1~[#7]~[#6]~[#6]~2~[#6](~[#6]~[#6]~3~[#6]~[#6]~[#6]~[#6,#7,#8,#16]~3)~[#6]~[#7]~[#6]~2~[#7]~1 |
| | | [#7]-[#6]~1~[#7]~[#6]~[#6]~2~[#6](~[#7]~[#6]~[#6]~2~[#6]~2~[#6]~[#6]~[#6]~[#6]~[#6]~2)~[#7]~1 |
| | | [#7]-[#6]~1~[#7]~[#6]~[#6]~2~[#6](~[#6]~[#6]~[#6]~3~[#6]~[#6]~[#6]~[#6,#7,#8,#16]~3)~[#6]~[#7]~[#6]~2~[#7]~1 |
| | | [#7]-[#6]~1~[#7]~[#6]~[#6]~2~[#6](~[#6]~[#6]~3~[#6]~[#6]~[#6]~[#6]~[#6]~3)~[#6]~[#7]~[#6]~2~[#7]~1 |
| | | [#7]-[#6]~1~[#7]~[#6]~[#6]~2~[#6,#7]~[#6](~[#6]~[#6]~3~[#6]~[#6]~[#6]~[#6,#7,#8,#16]~3)~[#6]~[#6,#7]~[#6]~2~[#7]~1 |
| | | [#7]-[#6]~1~[#7]~[#6]~[#6]~2~[#6,#7]~[#6](~[#6]~[#6,#7]~[#6]~2~[#7]~1)~[#6]~1~[#6]~[#6]~[#6]~[#6]~[#6]~1 |
| | | [#7]-[#6]~1~[#7]~[#6]~[#6]~2~[#6](~[#6]~[#6]~[#6]~3~[#6]~[#6]~[#6]~[#6]~[#6]~3)~[#6]~[#7]~[#6]~2~[#7]~1 |
| | | [#7]-[#6]~1~[#7]~[#6]~[#6]~2~[#6,#7]~[#6](~[#6]~[#6]~[#6]~3~[#6]~[#6]~[#6]~[#6,#7,#8,#16]~3)~[#6]~[#6,#7]~[#6]~2~[#7]~1 |
| | | [#7]-[#6]~1~[#7]~[#6]~[#6]~2~[#6,#7]~[#6](~[#6]~[#6]~3~[#6]~[#6]~[#6]~[#6]~[#6]~3)~[#6]~[#6,#7]~[#6]~2~[#7]~1 |
| | | [#7]-[#6]~1~[#7]~[#6]~[#6]~2~[#6,#7]~[#6](~[#6]~[#6]~[#6]~3~[#6]~[#6]~[#6]~[#6]~[#6]~3)~[#6]~[#6,#7]~[#6]~2~[#7]~1 |
| | | [#7]-*:1:*:*:*:*:2-*(-,=*-,=*-*:1:2)-[#6]-,=1-,=[#6]-,=[#6]-,=[#6]-,=[#8,#16]-,=1 |

| | | |
|---|---|---|
| | | [#7]-*:1:*:*:*:*:2-*(-,=*-,=*-*:1:2)-[#6,#7]-[#6]-,=1-,=[#6]-,=[#6]-,=[#6]-,=[#8,#16]-,=1 |
| | | [#7,#8]-,=*:1:*:*(-[#7]):*:*:2:*:*:*(:*:*:1:2)-c1ccccc1 |
| | | [#7,#8]-,=*:1:*:*(-[#7]):*:*:2:*:*:*(-[#6]-[#6,#7]-c3ccccc3):*:*:1:2 |
| mitotic inhibitors | | |
| | Taxane | |
| | | [#6]C1([#6])[#6]-2-[#6]-[#6]-[#6]=[#6]1-[#6]-[#6]-[#6]-[#6]-[#6]-2 |
| | Vinca_alkaloid | |
| | | [#6]~1~[#6]~[#6]~2~[#6]~[#7](~[#6]~1)~[#6]~[#6]~[#6]~1~[#6](~[#6]~[#6]~2)~[#7]~c2ccccc~12 |
| | | [#6]~1~[#6]~[#6]~2~[#6]~3~[#6]~[#6]~1~[#6]~[#7]~2~[#6]~[#6]~[#6]~1~[#6]~3~[#7]~c2ccccc~12 |
| | | [#6]~1~[#6]~[#6]~2~[#6]~[#6]~[#7]~3~[#6]~4~[#6](~[#6]~[#6]~[#7](~[#6]~1)~[#6]~2~4)~c1ccccc~31 |
| | | [#6]~1~[#6]~C~2~3~[#6](~[#6]~[#6]~[#6]~4~[#6]~[#6]~[#6]~[#7]~1~[#6]~2~4)~[#7]~c1ccccc~31 |
| topoisosmerase/gyrase inhibitors | | |
| | Camptothecin | |
| | | [#6]-1-[#8]-c2cc3-[#6]-[#6]-4-[#6]-[#8]-[#6]-[#6]-4-[#6]-c3cc2-[#8]-1 |
| | | [#8]-,=[#6]~1~[#6]~[#6]~[#6]~[#6]-2~[#7]~1-[#6]-c1cc3ccccc3nc-21 |
| | Antigyrase | |
| | | [#8]-,=[#6]~1~[#6](~[#6]~[#7](~c2cc(-[#7]-3-[#6]-[#6]-[$([#1,*])]-[#6]-[#6]-3)c(F)cc~12)-[#6]-1-[#6]-[#6]-1)-[#6](-[#8])=O' |
| | | [#8]-,=[#6]~1~[#6](~[#6]~[#7](-[#6]-[$([#1,*])])~c2nc(-[$([#1,*])])ccc~12)-[#6](-[#8])=O |

Supplementary Table 1 | Mapping of the different disease annotations between the Cellosaurus, Cell Model Passports and the Cancer Cell Line Encyclopedia (CCLE) databases

Cases of ambigous or inconsistent annotations between the different databases were manually curated.

| Cellosaurus | CCLE primary | CCLE subtype | CCLE primary + subtype | Cell Model Passports |
|---|---|---|---|---|
| Acute erythroid leukemia | Leukemia | Acute Myelogenous Leukemia (AML) | Leukemia - Acute Myelogenous Leukemia (AML) | Acute Myeloid Leukemia |
| Acute megakaryoblastic leukemia | Leukemia | Acute Myelogenous Leukemia (AML) | Leukemia - Acute Myelogenous Leukemia (AML) | Acute Myeloid Leukemia |
| Acute megakaryoblastic leukemia in Down syndrome | Leukemia | Acute Myelogenous Leukemia (AML) | Leukemia - Acute Myelogenous Leukemia (AML) | Acute Myeloid Leukemia |
| Acute monoblastic/monocytic leukemia | Leukemia | Acute Myelogenous Leukemia (AML) | Leukemia - Acute Myelogenous Leukemia (AML) | Acute Myeloid Leukemia |
| Acute myeloblastic leukemia with maturation | Leukemia | Acute Myelogenous Leukemia (AML) | Leukemia - Acute Myelogenous Leukemia (AML) | Acute Myeloid Leukemia |
| Acute myeloid leukemia | Leukemia | Acute Myelogenous Leukemia (AML) | Leukemia - Acute Myelogenous Leukemia (AML) | Acute Myeloid Leukemia |
| Acute myelomonocytic leukemia | Leukemia | Acute Myelogenous Leukemia (AML) | Leukemia - Acute Myelogenous Leukemia (AML) | Acute Myeloid Leukemia |
| Acute promyelocytic leukemia | Leukemia | Acute Myelogenous Leukemia (AML) | Leukemia - Acute Myelogenous Leukemia (AML) | Acute Myeloid Leukemia |
| Adenocarcinoma of ovary | Ovarian Cancer | Adenocarcinoma | Ovarian Cancer - Adenocarcinoma | Ovarian Carcinoma |
| Adenocarcinoma of the esophagus | Esophageal Cancer | Adenocarcinoma | Esophageal Cancer - Adenocarcinoma | Esophageal Carcinoma |
| Adenocarcinoma of the rat mammary gland | Breast Cancer | Adenocarcinoma | Breast Cancer - Adenocarcinoma | Breast Carcinoma |
| Adenocarcinoma of the rat prostate | Prostate Cancer | Adenocarcinoma | Prostate Cancer - Adenocarcinoma | Prostate Carcinoma |
| Adrenocortical carcinoma | Adrenal Cancer | Carcinoma | Adrenal Cancer - Carcinoma | Other Solid Cancers |
| Adult hepatocellular carcinoma | Liver Cancer | Hepatocellular Carcinoma | Liver Cancer - Hepatocellular Carcinoma | Hepatocellular Carcinoma |
| ALK-negative anaplastic large cell lymphoma | Lymphoma | T-cell | Lymphoma - T-cell | T-Cell Non-Hodgkin's Lymphoma |
| ALK-positive anaplastic large cell lymphoma | Lymphoma | T-cell | Lymphoma - T-cell | T-Cell Non-Hodgkin's Lymphoma |
| Alveolar rhabdomyosarcoma | Sarcoma | Rhabdomyosarcoma | Sarcoma - Rhabdomyosarcoma | Rhabdomyosarcoma |
| Amelanotic melanoma | Skin Cancer | Melanoma | Skin Cancer - Melanoma | Melanoma |
| Anaplastic astrocytoma | Brain Cancer | Astrocytoma | Brain Cancer - Astrocytoma | Glioma |
| Anaplastic thyroid carcinoma | Thyroid Cancer | Carcinoma | Thyroid Cancer - Carcinoma | Thyroid Gland Carcinoma |
| Astrocytoma | Brain Cancer | Astrocytoma | Brain Cancer - Astrocytoma | Glioma |
| B-cell chronic lymphocytic leukemia | Leukemia | Chronic Lymphoblastic Leukemia (CLL) | Leukemia - Chronic Lymphoblastic Leukemia (CLL) | Other Blood Cancers |
| B-cell non-Hodgkin lymphoma | Lymphoma | B-cell | Lymphoma - B-cell | B-Cell Non-Hodgkin's Lymphoma |
| Bladder carcinoma | Bladder Cancer | Carcinoma | Bladder Cancer - Carcinoma | Bladder Carcinoma |
| Bladder squamous cell carcinoma | Bladder Cancer | Squamous Cell Carcinoma | Bladder Cancer - Squamous Cell Carcinoma | Bladder Carcinoma |

| | | | | |
|---|---|---|---|---|
| Breast adenocarcinoma | Breast Cancer | Adenocarcinoma | Breast Cancer - Adenocarcinoma | Breast Carcinoma |
| Breast carcinoma | Breast Cancer | Carcinoma | Breast Cancer - Carcinoma | Breast Carcinoma |
| Breast ductal carcinoma | Breast Cancer | Breast Ductal Carcinoma | Breast Cancer - Breast Ductal Carcinoma | Breast Carcinoma |
| Breast inflammatory carcinoma | Breast Cancer | Breast Ductal Carcinoma | Breast Cancer - Breast Ductal Carcinoma | Breast Carcinoma |
| Breast pleomorphic carcinoma | Breast Cancer | Carcinoma | Breast Cancer - Carcinoma | Breast Carcinoma |
| Breast squamous cell carcinoma | Breast Cancer | Breast Ductal Carcinoma | Breast Cancer - Breast Ductal Carcinoma | Breast Carcinoma |
| Breast squamous cell carcinoma - acantholytic variant | Breast Cancer | Breast Ductal Carcinoma | Breast Cancer - Breast Ductal Carcinoma | Breast Carcinoma |
| Bronchogenic carcinoma | Lung Cancer | Non-Small Cell Lung Cancer (NSCLC) | Lung Cancer - Non-Small Cell Lung Cancer (NSCLC) | Other Solid Cancers |
| Burkitt lymphoma | Lymphoma | B-cell | Lymphoma - B-cell | Burkitt's Lymphoma |
| Canine soft tissue sarcoma | Sarcoma | Sarcoma | Sarcoma - Sarcoma | Other Sarcomas |
| Carcinoma of gallbladder and extrahepatic biliary tract | Gallbladder Cancer | Adenocarcinoma | Gallbladder Cancer - Adenocarcinoma | Biliary Tract Carcinoma |
| Carcinoma of the mouse prostate gland | Prostate Cancer | Adenocarcinoma | Prostate Cancer - Adenocarcinoma | Prostate Carcinoma |
| Cecum adenocarcinoma | Colon/Colorectal Cancer | Caecum Adenocarcinoma | Colon/Colorectal Cancer - Caecum Adenocarcinoma | Colorectal Carcinoma |
| Cervical carcinoma | Cervical Cancer | Carcinoma | Cervical Cancer - Carcinoma | Cervical Carcinoma |
| Cervical small cell carcinoma | Cervical Cancer | Small Cell Carcinoma | Cervical Cancer - Small Cell Carcinoma | Other Solid Cancers |
| Childhood T lymphoblastic lymphoma | Leukemia | Acute Lymphoblastic Leukemia (ALL) | Leukemia - Acute Lymphoblastic Leukemia (ALL) | T-Lymphoblastic Leukemia |
| Cholangiocarcinoma | Bile Duct Cancer | Cholangiocarcinoma | Bile Duct Cancer - Cholangiocarcinoma | Biliary Tract Carcinoma |
| Chondrosarcoma | Bone Cancer | Chondrosarcoma | Bone Cancer - Chondrosarcoma | Chondrosarcoma |
| Chronic eosinophilic leukemia | Leukemia | Acute Myelogenous Leukemia (AML) | Leukemia - Acute Myelogenous Leukemia (AML) | Acute Myeloid Leukemia |
| Chronic myeloid leukemia | Leukemia | Chronic Myelogenous Leukemia (CML) | Leukemia - Chronic Myelogenous Leukemia (CML) | Chronic Myelogenous Leukemia |
| Classic hairy cell leukemia | Leukemia | Hairy Cell | Leukemia - Hairy Cell | Other Blood Cancers |
| Clear cell adenocarcinoma of the ovary | Ovarian Cancer | Adenocarcinoma | Ovarian Cancer - Adenocarcinoma | Ovarian Carcinoma |
| Clear cell renal carcinoma | Kidney Cancer | Renal Carcinoma | Kidney Cancer - Renal Carcinoma | Kidney Carcinoma |
| Colon adenocarcinoma | Colon/Colorectal Cancer | Colon Adenocarcinoma | Colon/Colorectal Cancer - Colon Adenocarcinoma | Colorectal Carcinoma |
| Colon carcinoma | Colon/Colorectal Cancer | Carcinoma | Colon/Colorectal Cancer - Carcinoma | Colorectal Carcinoma |
| Colorectal adenocarcinoma | Colon/Colorectal Cancer | Adenocarcinoma | Colon/Colorectal Cancer - Adenocarcinoma | Colorectal Carcinoma |
| Colorectal carcinoma | Colon/Colorectal Cancer | Carcinoma | Colon/Colorectal Cancer - Carcinoma | Colorectal Carcinoma |
| Cutaneous melanoma | Skin Cancer | Melanoma | Skin Cancer - Melanoma | Melanoma |

| | | | | |
|---|---|---|---|---|
| Cystic fibrosis | Pancreatic Cancer | Ductal Adenocarcinoma | Pancreatic Cancer - Ductal Adenocarcinoma | Pancreatic Carcinoma |
| Differentiated thyroid carcinoma | Thyroid Cancer | Carcinoma | Thyroid Cancer - Carcinoma | Thyroid Gland Carcinoma |
| Diffuse large B-cell lymphoma | Lymphoma | Diffuse Large B-cell Lymphoma (DLBCL) | Lymphoma - Diffuse Large B-cell Lymphoma (DLBCL) | B-Cell Non-Hodgkin's Lymphoma |
| Duodenal adenocarcinoma | Gastric Cancer | Duodenal Adenocarcinoma | Gastric Cancer - Duodenal Adenocarcinoma | Other Solid Cancers |
| Embryonal carcinoma | Embryonal Cancer | Carcinoma | Embryonal Cancer - Carcinoma | Other Solid Cancers |
| Embryonal rhabdomyosarcoma | Rhabdoid | Rhabdomyosarcoma | Sarcoma - Rhabdomyosarcoma | Rhabdomyosarcoma |
| Endometrial adenocarcinoma | Endometrial/Uterine Cancer | Endometrial Adenocarcinoma | Endometrial/Uterine Cancer - Endometrial Adenocarcinoma | Endometrial Carcinoma |
| Endometrial adenosquamous carcinoma | Endometrial/Uterine Cancer | Endometrial Adenosquamous Carcinoma | Endometrial/Uterine Cancer - Endometrial Adenosquamous Carcinoma | Endometrial Carcinoma |
| Endometrial carcinoma | Endometrial/Uterine Cancer | Endometrial Adenocarcinoma | Endometrial/Uterine Cancer - Endometrial Adenocarcinoma | Endometrial Carcinoma |
| Endometrial stromal sarcoma | Endometrial/Uterine Cancer | Endometrial Stromal Sarcoma | Endometrial/Uterine Cancer - Endometrial Stromal Sarcoma | Other Sarcomas |
| Endometrioid carcinoma of ovary | Ovarian Cancer | Adenocarcinoma | Ovarian Cancer - Adenocarcinoma | Ovarian Carcinoma |
| Epithelioid sarcoma | Sarcoma | Epithelioid | Sarcoma - Epithelioid | Other Sarcomas |
| Ewing sarcoma | Bone Cancer | Ewings Sarcoma | Bone Cancer - Ewings Sarcoma | Ewing's Sarcoma |
| Extraskeletal myxoid chondrosarcoma | Bone Cancer | Chondrosarcoma | Bone Cancer - Chondrosarcoma | Chondrosarcoma |
| Fibrosarcoma | Sarcoma | Fibrosarcoma | Sarcoma - Fibrosarcoma | Other Sarcomas |
| Gastric adenocarcinoma | Gastric Cancer | Adenocarcinoma | Gastric Cancer - Adenocarcinoma | Gastric Carcinoma |
| Gastric adenosquamous carcinoma | Gastric Cancer | Adenocarcinoma | Gastric Cancer - Adenocarcinoma | Gastric Carcinoma |
| Gastric carcinoma | Gastric Cancer | Adenocarcinoma | Gastric Cancer - Adenocarcinoma | Gastric Carcinoma |
| Gastric choriocarcinoma | Gastric Cancer | Choriocarcinoma | Gastric Cancer - Choriocarcinoma | Gastric Carcinoma |
| Gastric signet ring cell adenocarcinoma | Gastric Cancer | Adenocarcinoma | Gastric Cancer - Adenocarcinoma | Gastric Carcinoma |
| Gastric small cell carcinoma | Gastric Cancer | Small Cell Carcinoma | Gastric Cancer - Small Cell Carcinoma | Gastric Carcinoma |
| Gastric tubular adenocarcinoma | Gastric Cancer | Adenocarcinoma | Gastric Cancer - Adenocarcinoma | Gastric Carcinoma |
| Gastrointestinal stromal tumor | Gastric Cancer | Carcinoma | Gastric Cancer – Carcinoma | Gastric Carcinoma |
| Gestational choriocarcinoma | Endometrial/Uterine Cancer | Choriocarcinoma | Endometrial/Uterine Cancer - Choriocarcinoma | Other Solid Cancers |
| Glioblastoma | Brain Cancer | Glioblastoma | Brain Cancer - Glioblastoma | Glioblastoma |
| Gliosarcoma | Brain Cancer | Glioma | Brain Cancer - Glioma | Glioblastoma |
| Head and neck squamous cell carcinoma | Head and Neck Cancer | Squamous Cell Carcinoma | Head and Neck Cancer - Squamous Cell Carcinoma | Head and Neck Carcinoma |
| Hepatoblastoma | Liver Cancer | Hepatoblastoma | Liver Cancer - Hepatoblastoma | Hepatocellular Carcinoma |

| | | | | |
|---|---|---|---|---|
| Hepatocellular carcinoma | Liver Cancer | Hepatocellular Carcinoma | Liver Cancer - Hepatocellular Carcinoma | Hepatocellular Carcinoma |
| Hepatocellular carcinoma of the mouse | Liver Cancer | Hepatocellular Carcinoma | Liver Cancer - Hepatocellular Carcinoma | Hepatocellular Carcinoma |
| High grade ovarian serous adenocarcinoma | Ovarian Cancer | Adenocarcinoma | Ovarian Cancer - Adenocarcinoma | Ovarian Carcinoma |
| Hodgkin lymphoma | Lymphoma | B-cell | Lymphoma - B-cell | Hodgkin's Lymphoma |
| Human papillomavirus-related endocervical adenocarcinoma | Cervical Cancer | Cervical Adenocarcinoma | Cervical Cancer - Cervical Adenocarcinoma | Cervical Carcinoma |
| Invasive breast carcinoma of no special type | Breast Cancer | Carcinoma | Breast Cancer - Carcinoma | Breast Carcinoma |
| Invasive breast lobular carcinoma | Breast Cancer | Breast Ductal Carcinoma | Breast Cancer - Breast Ductal Carcinoma | Breast Carcinoma |
| Kidney neoplasm | Kidney Cancer | Renal Leiomyoblastoma | Kidney Cancer - Renal Leiomyoblastoma | Other Sarcomas |
| Leiomyosarcoma of the corpus uteri | Sarcoma | Leiomyosarcoma | Sarcoma - Leiomyosarcoma | Other Sarcomas |
| Liposarcoma | Liposarcoma | Sarcoma | Liposarcoma - Sarcoma | Other Sarcomas |
| Lung adenocarcinoma | Lung Cancer | Non-Small Cell Lung Cancer (NSCLC) | Lung Cancer - Non-Small Cell Lung Cancer (NSCLC) | Non-Small Cell Lung Carcinoma |
| Lung adenosquamous carcinoma | Lung Cancer | Non-Small Cell Lung Cancer (NSCLC) | Lung Cancer - Non-Small Cell Lung Cancer (NSCLC) | Non-Small Cell Lung Carcinoma |
| Lung carcinoid tumor | Lung Cancer | Carcinoid | Lung Cancer - Carcinoid | Other Solid Cancers |
| Lung carcinoma | Lung Cancer | Carcinoma | Lung Cancer - Carcinoma | Other Solid Cancers |
| Lung giant cell carcinoma | Lung Cancer | Non-Small Cell Lung Cancer (NSCLC) | Lung Cancer - Non-Small Cell Lung Cancer (NSCLC) | Non-Small Cell Lung Carcinoma |
| Lung large cell carcinoma | Lung Cancer | Non-Small Cell Lung Cancer (NSCLC) | Lung Cancer - Non-Small Cell Lung Cancer (NSCLC) | Non-Small Cell Lung Carcinoma |
| Lung mucoepidermoid carcinoma | Lung Cancer | Non-Small Cell Lung Cancer (NSCLC) | Lung Cancer - Non-Small Cell Lung Cancer (NSCLC) | Other Solid Cancers |
| Lung non-small cell carcinoma | Lung Cancer | Non-Small Cell Lung Cancer (NSCLC) | Lung Cancer - Non-Small Cell Lung Cancer (NSCLC) | Non-Small Cell Lung Carcinoma |
| Lung papillary adenocarcinoma | Lung Cancer | Non-Small Cell Lung Cancer (NSCLC) | Lung Cancer - Non-Small Cell Lung Cancer (NSCLC) | Non-Small Cell Lung Carcinoma |
| Lung squamous cell carcinoma | Lung Cancer | Non-Small Cell Lung Cancer (NSCLC) | Lung Cancer - Non-Small Cell Lung Cancer (NSCLC) | Non-Small Cell Lung Carcinoma |
| Lymphoblastic lymphoma | Lymphoma | Lymphoblastic Lymphoma | Lymphoma - Lymphoblastic Lymphoma | B-Cell Non-Hodgkin's Lymphoma |
| Malignant mixed Mullerian tumor of the ovary | Ovarian Cancer | Carcinosarcoma | Ovarian Cancer - Carcinosarcoma | Ovarian Carcinoma |
| Malignant neoplasms of the mouse mammary gland | Breast Cancer | Adenocarcinoma | Breast Cancer - Adenocarcinoma | Breast Carcinoma |
| Malignant tumors of the mouse pulmonary system | Lung Cancer | Carcinoma | Lung Cancer - Carcinoma | Other Solid Cancers |
| Mantle cell lymphoma | Lymphoma | B-cell | Lymphoma - B-cell | B-Cell Non-Hodgkin's Lymphoma |
| Mast cell leukemia | Leukemia | Multiple Myeloma | Leukemia - Multiple Myeloma | Other Blood Cancers |
| Medulloblastoma | Brain Cancer | Medulloblastoma | Brain Cancer - Medulloblastoma | Other Solid Cancers |
| Melanoma | Skin Cancer | Melanoma | Skin Cancer - Melanoma | Melanoma |

| | | | | |
|---|---|---|---|---|
| Minimally invasive lung adenocarcinoma | Lung Cancer | Non-Small Cell Lung Cancer (NSCLC) | Lung Cancer - Non-Small Cell Lung Cancer (NSCLC) | Non-Small Cell Lung Carcinoma |
| Mixed germ cell tumor | Ovarian Cancer | Carcinosarcoma | Ovarian Cancer - Carcinosarcoma | Ovarian Carcinoma |
| Mouse colon adenocarcinoma | Colon/Colorectal Cancer | Adenocarcinoma | Colon/Colorectal Cancer - Adenocarcinoma | Colorectal Carcinoma |
| Mouse fibrosarcoma | Fibroblast | Fibrosarcoma | Fibroblast - Fibrosarcoma | Other Sarcomas |
| Mouse glioblastoma | Brain Cancer | Glioblastoma | Brain Cancer - Glioblastoma | Glioblastoma |
| Mouse kidney carcinoma | Kidney Cancer | Carcinoma | Kidney Cancer - Carcinoma | Other Solid Cancers |
| Mouse leukemia | Leukemia | Leukemia | Leukemia - Leukemia | Other Blood Cancers |
| Mouse Leydig cell tumor | Testicular Cancer | Leydig cell | Testicular Cancer - Leydig cell | Testicular Carcinoma |
| Mouse lymphoma | Lymphoma | Lymphoma | Lymphoma - Lymphoma | Other Blood Cancers |
| Mouse mast cell neoplasm | Myeloma | Multiple Myeloma | Myeloma - Multiple Myeloma | Other Blood Cancers |
| Mouse melanoma | Skin Cancer | Melanoma | Skin Cancer - Melanoma | Melanoma |
| Mouse neuroblastoma | Neuroblastoma | Neuroblastoma | Neuroblastoma – Neuroblastoma | Neuroblastoma |
| Mouse pancreatic ductal adenocarcinoma | Pancreatic Cancer | Ductal Adenocarcinoma | Pancreatic Cancer - Ductal Adenocarcinoma | Pancreatic Carcinoma |
| Mouse plasma cell myeloma | Myeloma | Plasma cell | Myeloma - Plasma cell | Plasma Cell Myeloma |
| Mouse reticulum cell sarcoma | Sarcoma | Fibrosarcoma | Sarcoma - Fibrosarcoma | Other Solid Cancers |
| Mouse sarcoma | Sarcoma | Sarcoma | Sarcoma – Sarcoma | Other Sarcomas |
| Mouse squamous cell carcinoma | Skin Cancer | Squamous Cell Carcinoma | Skin Cancer - Squamous Cell Carcinoma | Squamous Cell Lung Carcinoma |
| Mouse teratocarcinoma | Liposarcoma | Sarcoma | Liposarcoma - Sarcoma | Other Sarcomas |
| Mouse thymic lymphoma | Lymphoma | B-cell | Lymphoma - B-cell | Other Blood Cancers |
| Mucinous adenocarcinoma of ovary | Ovarian Cancer | Adenocarcinoma | Ovarian Cancer - Adenocarcinoma | Ovarian Carcinoma |
| Multiple endocrine neoplasia type 2 | Thyroid Cancer | Squamous Cell Carcinoma | Thyroid Cancer - Squamous Cell Carcinoma | Thyroid Gland Carcinoma |
| Multiple myeloma | Myeloma | Multiple Myeloma | Myeloma - Multiple Myeloma | Plasma Cell Myeloma |
| Mycosis fungoides and Sezary syndrome | Lymphoma | T-cell | Lymphoma - T-cell | T-Cell Non-Hodgkin's Lymphoma |
| Natural killer cell lymphoblastic leukemia/lymphoma | Leukemia | Natural Killer Cell Lymphoblastic Leukemia/Lymphoma | Leukemia - Natural Killer Cell Lymphoblastic Leukemia/Lymphoma | Other Blood Cancers |
| Neuroblastoma | Neuroblastoma | Neuroblastoma | Neuroblastoma - Neuroblastoma | Neuroblastoma |
| Non-central nervous system-localized embryonal carcinoma | Embryonal Cancer | Carcinoma | Embryonal Cancer - Carcinoma | Other Solid Cancers |
| Oligodendroglioma | Brain Cancer | Oligodendroglioma | Brain Cancer - Oligodendroglioma | Glioma |
| Oral epithelial dysplasia | Head and Neck Cancer | Oral Dysplasia | Head and Neck Cancer - Oral Dysplasia | Oral Cavity Carcinoma |
| Osteosarcoma | Bone Cancer | Osteosarcoma | Bone Cancer - Osteosarcoma | Osteosarcoma |
| Ovarian carcinoma | Ovarian Cancer | Carcinoma | Ovarian Cancer - Carcinoma | Ovarian Carcinoma |
| Ovarian cystadenocarcinoma | Ovarian Cancer | Cystadenocarcinoma | Ovarian Cancer - Cystadenocarcinoma | Ovarian Carcinoma |

| | | | | |
|---|---|---|---|---|
| Ovarian granulosa cell tumor | Ovarian Cancer | Granulosa Cell Tumor | Ovarian Cancer - Granulosa Cell Tumor | Other Solid Cancers |
| Ovarian serous adenocarcinoma | Ovarian Cancer | Cystadenocarcinoma | Ovarian Cancer - Cystadenocarcinoma | Ovarian Carcinoma |
| Ovarian serous cystadenocarcinoma | Ovarian Cancer | Cystadenocarcinoma | Ovarian Cancer - Cystadenocarcinoma | Ovarian Carcinoma |
| Pancreatic adenocarcinoma | Pancreatic Cancer | Ductal Adenocarcinoma | Pancreatic Cancer - Ductal Adenocarcinoma | Pancreatic Carcinoma |
| Pancreatic carcinoma | Pancreatic Cancer | Ductal Adenocarcinoma | Pancreatic Cancer - Ductal Adenocarcinoma | Pancreatic Carcinoma |
| Pancreatic ductal adenocarcinoma | Pancreatic Cancer | Ductal Adenocarcinoma | Pancreatic Cancer - Ductal Adenocarcinoma | Pancreatic Carcinoma |
| Papillary renal cell carcinoma | Kidney Cancer | Renal Cell Carcinoma | Kidney Cancer - Renal Cell Carcinoma | Kidney Carcinoma |
| Papilloma of the mouse skin | Skin Cancer | Squamous Cell Carcinoma | Skin Cancer - Squamous Cell Carcinoma | Other Solid Cancers |
| Pediatric hepatocellular carcinoma | Liver Cancer | Hepatocellular Carcinoma | Liver Cancer - Hepatocellular Carcinoma | Hepatocellular Carcinoma |
| Pleural mesothelioma | Lung Cancer | Mesothelioma | Lung Cancer - Mesothelioma | Mesothelioma |
| Poorly differentiated thyroid gland carcinoma | Thyroid Cancer | Carcinoma | Thyroid Cancer - Carcinoma | Thyroid Gland Carcinoma |
| Precursor B-cell acute lymphoblastic leukemia | Leukemia | Acute Lymphoblastic Leukemia (ALL) | Leukemia - Acute Lymphoblastic Leukemia (ALL) | B-Lymphoblastic Leukemia |
| Precursor T-cell acute lymphoblastic leukemia | Leukemia | Acute Lymphoblastic Leukemia (ALL) | Leukemia - Acute Lymphoblastic Leukemia (ALL) | T-Lymphoblastic Leukemia |
| Primary cutaneous T-cell non-Hodgkin lymphoma | Lymphoma | T-cell | Lymphoma - T-cell | T-Cell Non-Hodgkin's Lymphoma |
| Primary effusion lymphoma | Lymphoma | B-cell | Lymphoma - B-cell | B-Cell Non-Hodgkin's Lymphoma |
| Primitive neuroectodermal tumor | Brain Cancer | Primitive Neuroectodermal Tumor (PNET) | Brain Cancer - Primitive Neuroectodermal Tumor (PNET) | Other Solid Cancers |
| Prostate carcinoma | Prostate Cancer | Adenocarcinoma | Prostate Cancer - Adenocarcinoma | Prostate Carcinoma |
| Rat adrenal gland pheochromocytoma | Adrenal Cancer | Adenocarcinoma | Adrenal Cancer - Adenocarcinoma | Other Solid Cancers |
| Rat digestive system neoplasms | Colon/Colorectal Cancer | Carcinoma | Colon/Colorectal Cancer – Carcinoma | Colorectal Carcinoma |
| Rat hepatocellular carcinoma | Liver Cancer | Hepatocellular Carcinoma | Liver Cancer - Hepatocellular Carcinoma | Hepatocellular Carcinoma |
| Rat insulinoma | Pancreatic Cancer | Adenocarcinoma | Pancreatic Cancer - Adenocarcinoma | Pancreatic Carcinoma |
| Rat leukemia | Leukemia | Leukemia | Leukemia - Leukemia | Other Blood Cancers |
| Rat malignant glioma | Brain Cancer | Glioma | Brain Cancer - Glioma | Glioma |
| Rat pituitary gland neoplasm | Brain Cancer | Glioma | Brain Cancer – Glioma | Glioma |
| Rat sarcoma | Sarcoma | Sarcoma | Sarcoma - Sarcoma | Other Sarcomas |
| Rectal adenocarcinoma | Colon/Colorectal Cancer | Adenocarcinoma | Colon/Colorectal Cancer - Adenocarcinoma | Colorectal Carcinoma |
| Recurrent bladder carcinoma | Bladder Cancer | Carcinoma | Bladder Cancer - Carcinoma | Bladder Carcinoma |
| Renal cell carcinoma | Kidney Cancer | Renal Cell Carcinoma | Kidney Cancer - Renal Cell Carcinoma | Kidney Carcinoma |

| | | | | |
|---|---|---|---|---|
| Renal pelvis and ureter urothelial carcinoma | Kidney Cancer | Renal Carcinoma | Kidney Cancer - Renal Carcinoma | Kidney Carcinoma |
| Retinoblastoma | Eye Cancer | Retinoblastoma | Eye Cancer - Retinoblastoma | Other Solid Cancers |
| Rhabdomyosarcoma | Sarcoma | Rhabdomyosarcoma | Sarcoma - Rhabdomyosarcoma | Rhabdomyosarcoma |
| Sarcoma of the corpus uteri | Sarcoma | Uterine Sarcoma | Sarcoma - Uterine Sarcoma | Other Sarcomas |
| Skin squamous cell carcinoma | Skin Cancer | Squamous Cell Carcinoma | Skin Cancer - Squamous Cell Carcinoma | Other Solid Cancers |
| Small cell lung cancer | Lung Cancer | Small Cell Lung Cancer (SCLC) | Lung Cancer - Small Cell Lung Cancer (SCLC) | Small Cell Lung Carcinoma |
| Squamous cell carcinoma of pancreas | Pancreatic Cancer | Squamous Cell Carcinoma | Pancreatic Cancer - Squamous Cell Carcinoma | Pancreatic Carcinoma |
| Squamous cell carcinoma of salivary glands | Head and Neck Cancer | Squamous Cell Carcinoma | Head and Neck Cancer - Squamous Cell Carcinoma | Head and Neck Carcinoma |
| Squamous cell carcinoma of the cervix uteri | Cervical Cancer | Squamous Cell Carcinoma | Cervical Cancer - Squamous Cell Carcinoma | Cervical Carcinoma |
| Squamous cell carcinoma of the esophagus | Esophageal Cancer | Squamous Cell Carcinoma | Esophageal Cancer - Squamous Cell Carcinoma | Esophageal Squamous Cell Carcinoma |
| Squamous cell carcinoma of the hypopharynx | Head and Neck Cancer | Squamous Cell Carcinoma | Head and Neck Cancer - Squamous Cell Carcinoma | Head and Neck Carcinoma |
| Squamous cell carcinoma of the larynx | Head and Neck Cancer | Squamous Cell Carcinoma | Head and Neck Cancer - Head and Neck Cancer | Head and Neck Carcinoma |
| Squamous cell carcinoma of the mouse skin | Head and Neck Cancer | Squamous Cell Carcinoma | Head and Neck Cancer - Head and Neck Cancer | Head and Neck Carcinoma |
| Squamous cell carcinoma of the oral cavity | Head and Neck Cancer | Squamous Cell Carcinoma | Head and Neck Cancer - Squamous Cell Carcinoma | Oral Cavity Carcinoma |
| Squamous cell carcinoma of the oral tongue | Head and Neck Cancer | Squamous Cell Carcinoma | Head and Neck Cancer - Squamous Cell Carcinoma | Oral Cavity Carcinoma |
| Synovial sarcoma | Sarcoma | Synovial | Sarcoma - Synovial | Other Sarcomas |
| Thyroid gland sarcoma | Sarcoma | Thyroid Sarcoma | Sarcoma - Thyroid Sarcoma | Thyroid Gland Carcinoma |
| Undifferentiated pleomorphic sarcoma | Sarcoma | Pleomorphic Sarcoma | Sarcoma - Pleomorphic Sarcoma | Other Sarcomas |
| Ureter urothelial carcinoma | Bladder Cancer | Transitional Cell Carcinoma | Bladder Cancer - Transitional Cell Carcinoma | Bladder Carcinoma |
| Vaginal melanoma | Skin Cancer | Melanoma | Skin Cancer - Melanoma | Melanoma |
| Vulvar carcinoma | Skin Cancer | Carcinoma | Skin Cancer - Carcinoma | Other Solid Cancers |
| Vulvar leiomyosarcoma | Sarcoma | Leiomyosarcoma | Sarcoma - Leiomyosarcoma | Other Sarcomas |
| Vulvar melanoma | Skin Cancer | Melanoma | Skin Cancer - Melanoma | Melanoma |
| Vulvar squamous cell carcinoma | Skin Cancer | Squamous Cell Carcinoma | Skin Cancer - Squamous Cell Carcinoma | Other Solid Cancers |

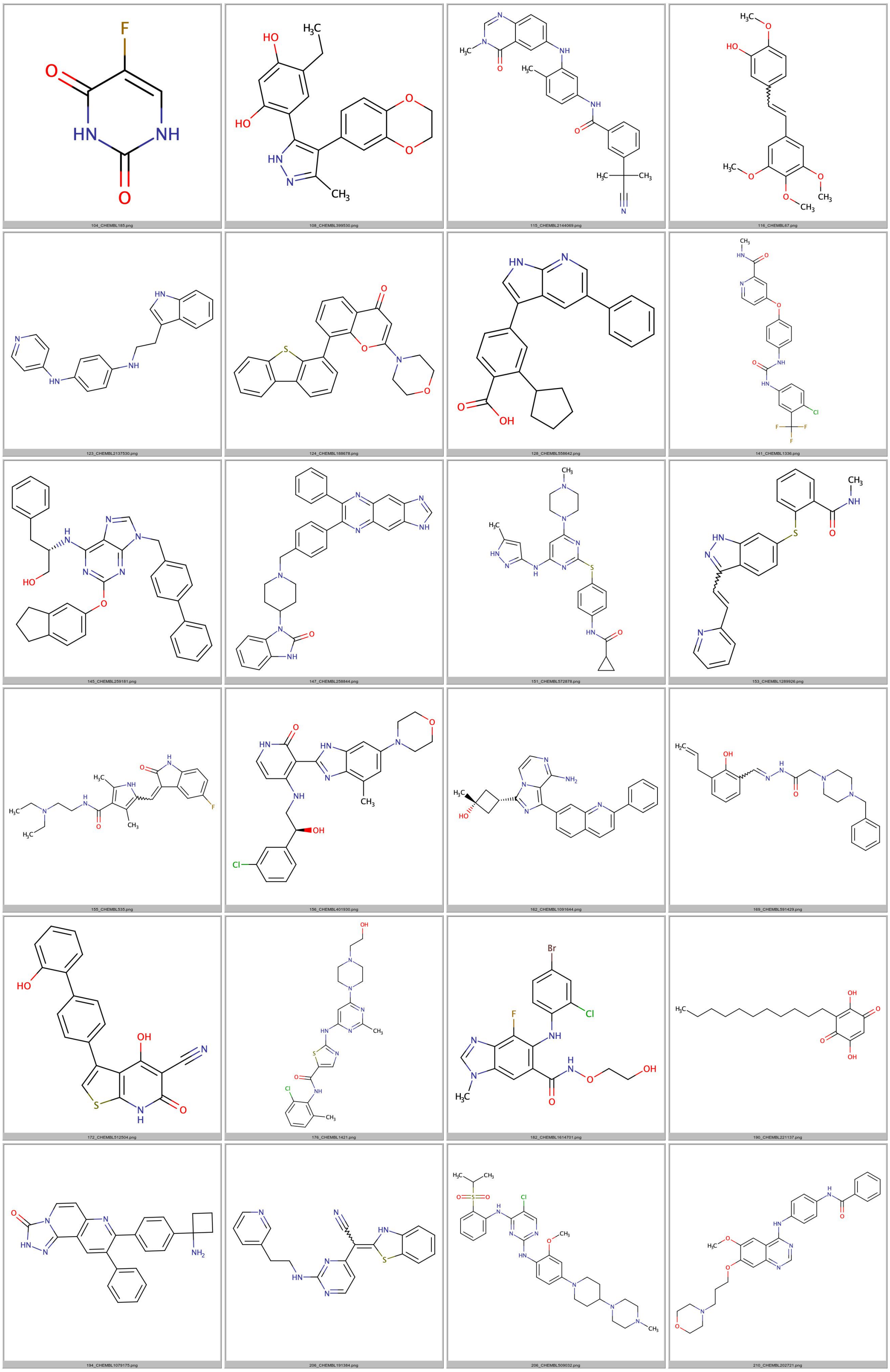

Supplementary Figure 1 | Chemical structures of the 79 objective ubiquitous compounds
Compounds were considered objective ubiquitous when active on more than 100 cell lines.

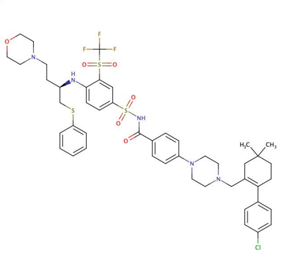

217_CHEMBL443684.png

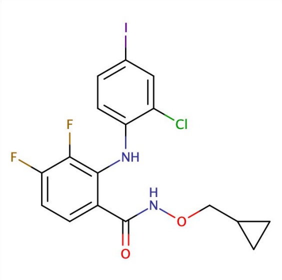

222_CHEMBL105442.png

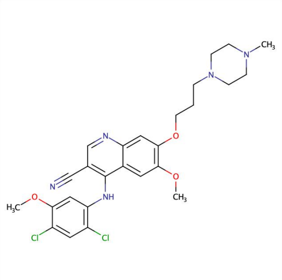

225_CHEMBL288441.png

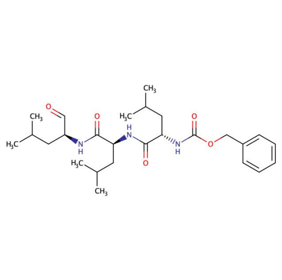

233_CHEMBL64925.png

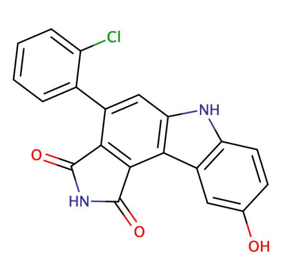

241_CHEMBL379975.png

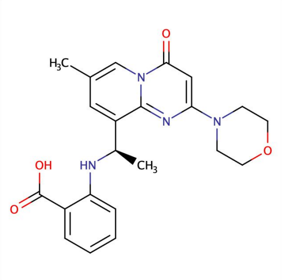

242_CHEMBL2165191.png

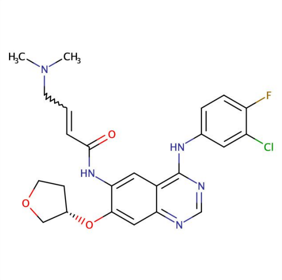

248_CHEMBL1173655.png

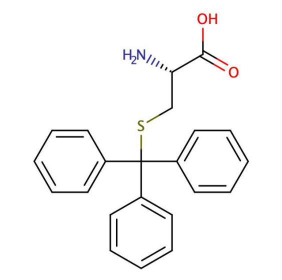

250_CHEMBL392695.png

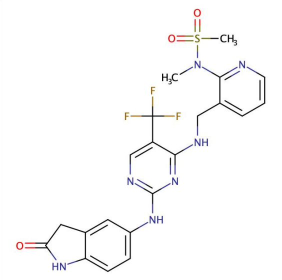

252_CHEMBL1084546.png

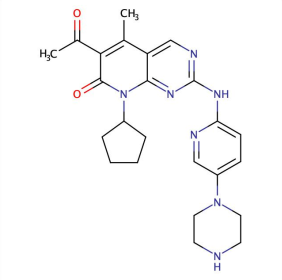

258_CHEMBL189963.png

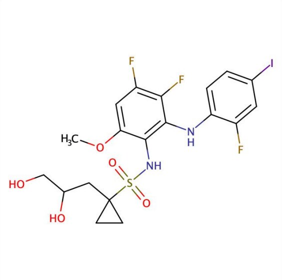

260_CHEMBL3349342.png

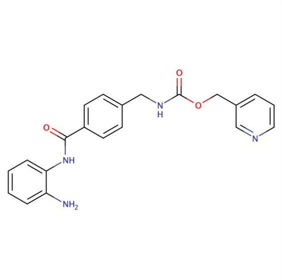

268_CHEMBL27759.png

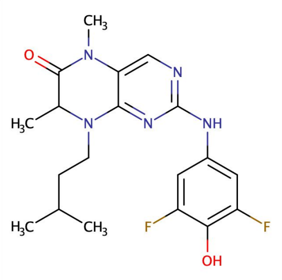

268_CHEMBL573107.png

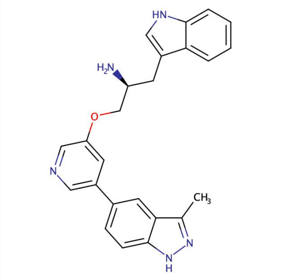

281_CHEMBL379300.png

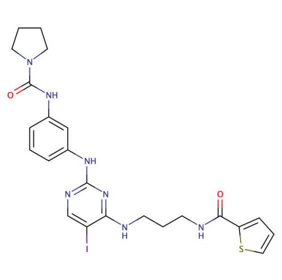

284_CHEMBL577784.png

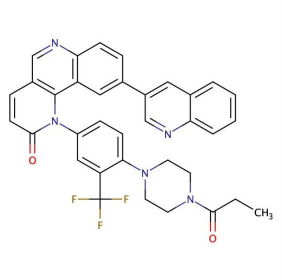

285_CHEMBL1256459.png

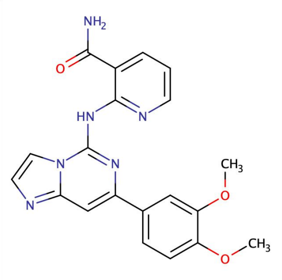

289_CHEMBL1190711.png

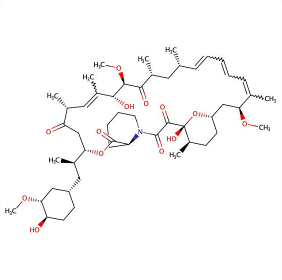

294_CHEMBL413.png

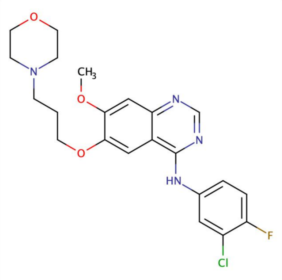

295_CHEMBL939.png

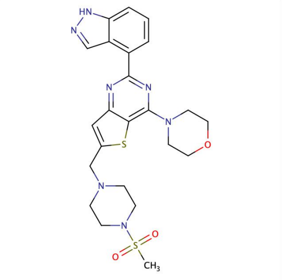

298_CHEMBL521851.png

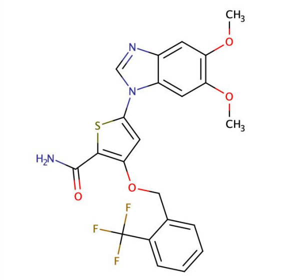

299_CHEMBL514499.png

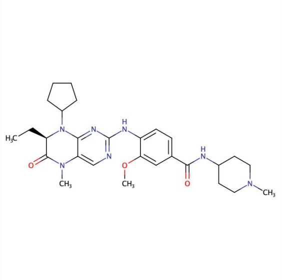

301_CHEMBL513909.png

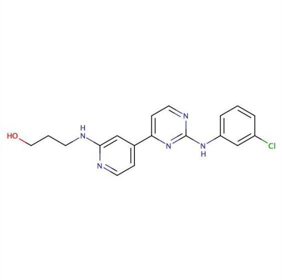

313_CHEMBL300389.png

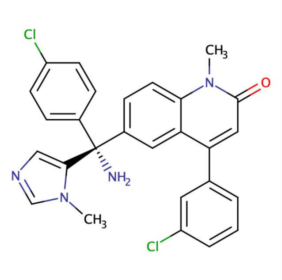

328_CHEMBL289228.png

Supplementary Figure 2 (continued)

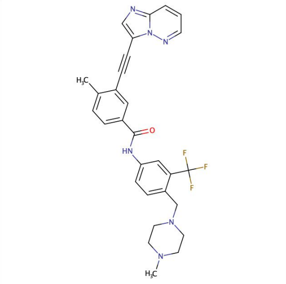
361_CHEMBL1171837.png

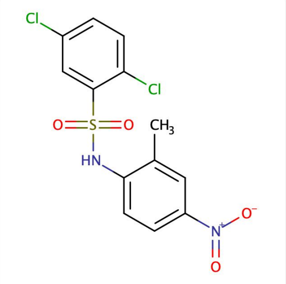
376_CHEMBL2363137.png

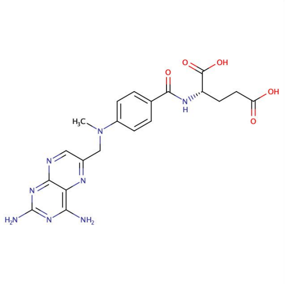
397_CHEMBL34259.png

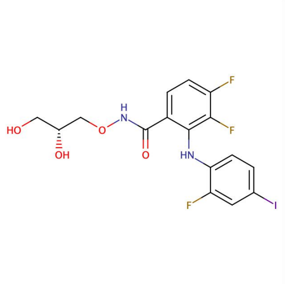
422_CHEMBL507361.png

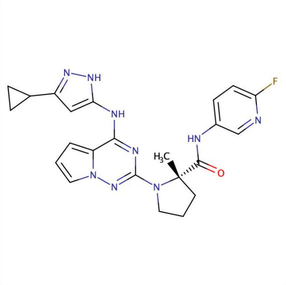
423_CHEMBL575448.png

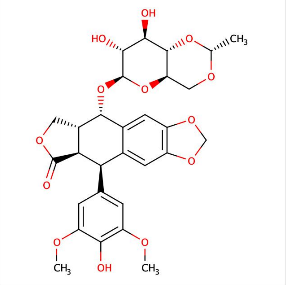
435_CHEMBL44657.png

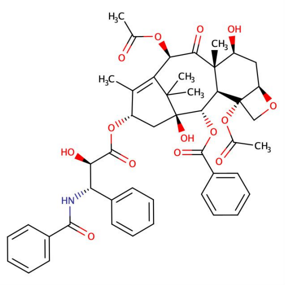
439_CHEMBL428647.png

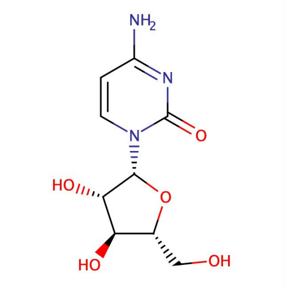
439_CHEMBL803.png

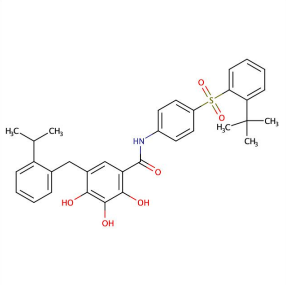
441_CHEMBL217354.png

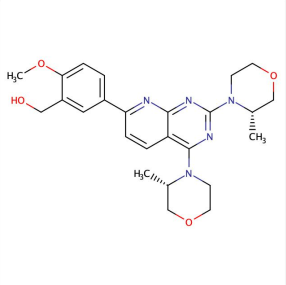
467_CHEMBL1801204.png

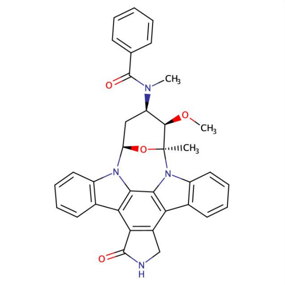
472_CHEMBL608533.png

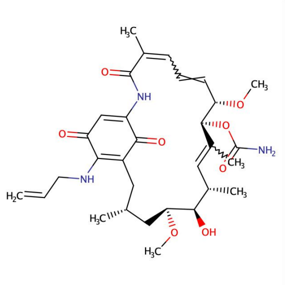
477_CHEMBL109480.png

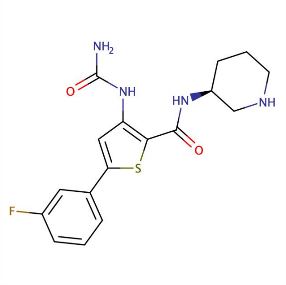
478_CHEMBL2041933.png

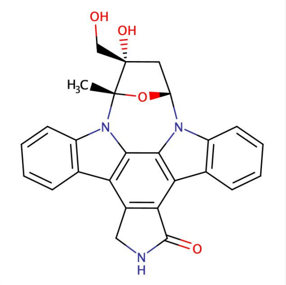
479_CHEMBL603469.png

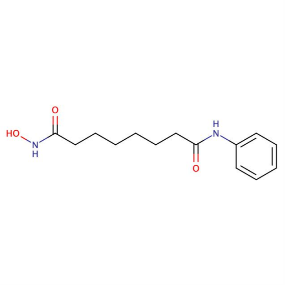
493_CHEMBL98.png

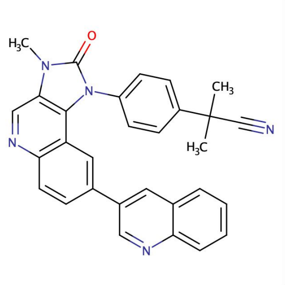
518_CHEMBL1879463.png

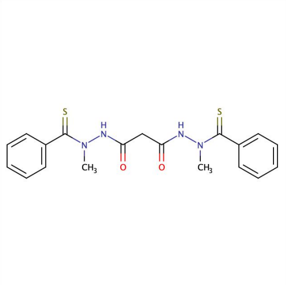
518_CHEMBL1972860.png

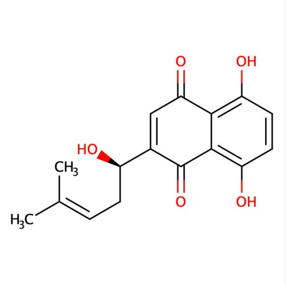
535_CHEMBL9470.png

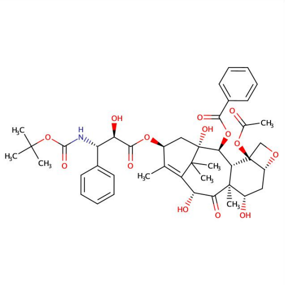
538_CHEMBL3545252.png

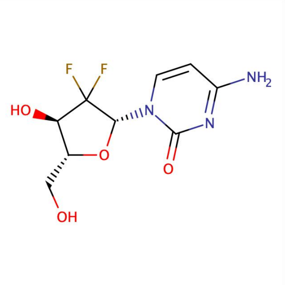
548_CHEMBL888.png

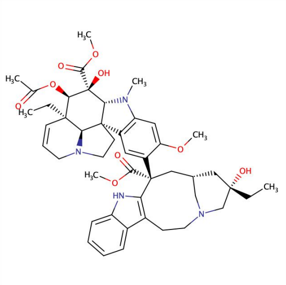
549_CHEMBL159.png

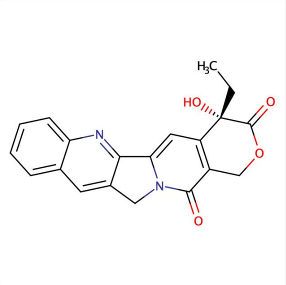
568_CHEMBL65.png

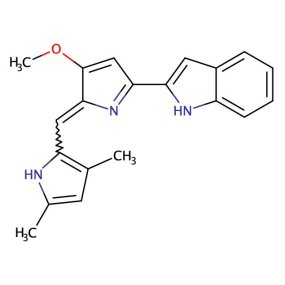
572_CHEMBL2107358.png

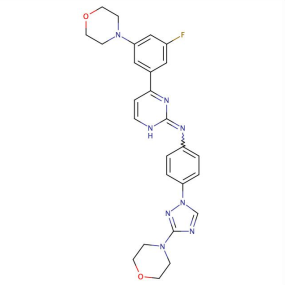
572_CHEMBL585951.png

Supplementary Figure 3 (continued)

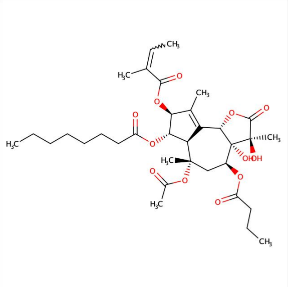
593_CHEMBL96926.png

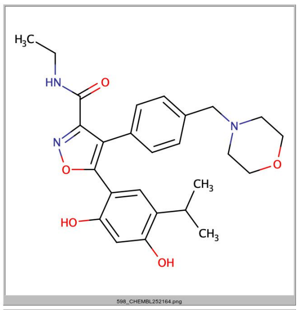
598_CHEMBL252164.png

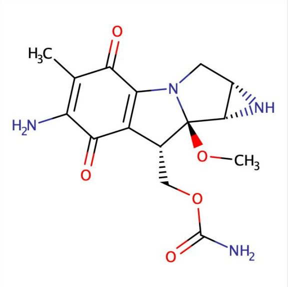
609_CHEMBL105.png

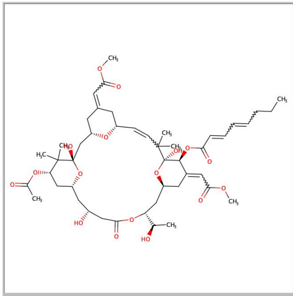
612_CHEMBL449158.png

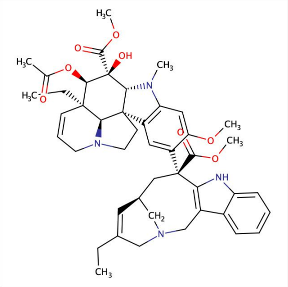
616_CHEMBL553025.png

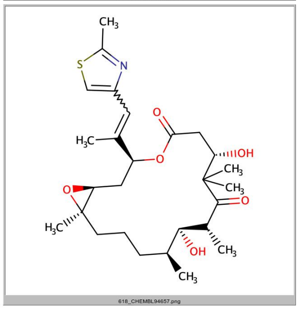
618_CHEMBL94657.png

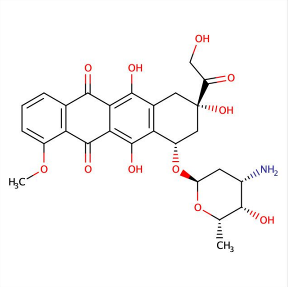
665_CHEMBL53463.png

Supplementary Figure 4 (continued)

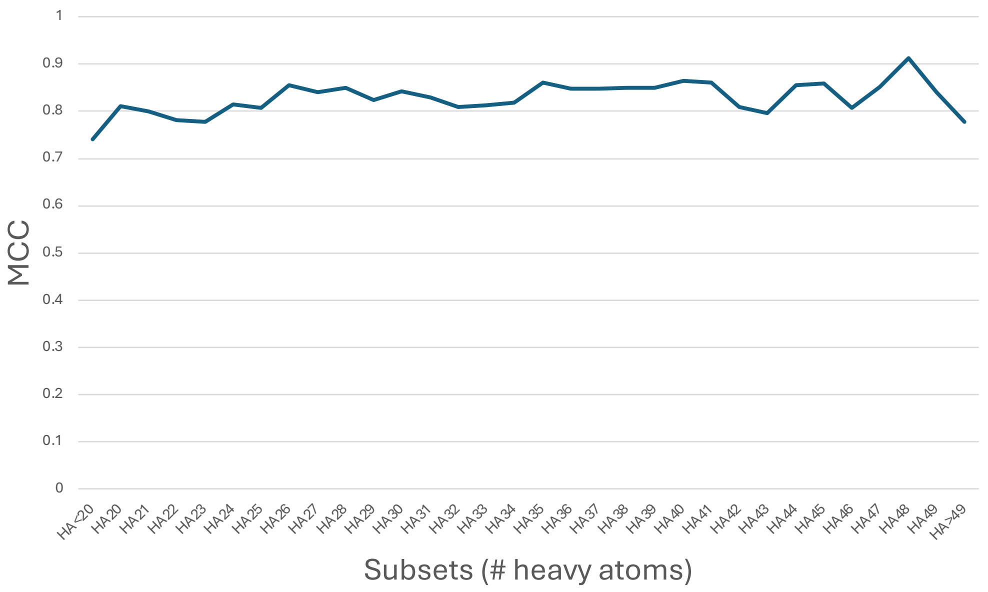

| Subset | MCC |
|---|---|
| HA<20 | 0.74076 |
| HA20 | 0.81107 |
| HA21 | 0.79988 |
| HA22 | 0.78184 |
| HA23 | 0.77867 |
| HA24 | 0.8155 |
| HA25 | 0.8076 |
| HA26 | 0.85488 |
| HA27 | 0.84128 |
| HA28 | 0.85044 |
| HA29 | 0.82394 |
| HA30 | 0.84153 |
| HA31 | 0.82975 |
| HA32 | 0.81006 |
| HA33 | 0.81245 |
| HA34 | 0.81933 |
| HA35 | 0.86166 |
| HA36 | 0.84799 |
| HA37 | 0.84728 |
| HA38 | 0.8504 |
| HA39 | 0.85021 |
| HA40 | 0.86398 |
| HA41 | 0.8605 |
| HA42 | 0.80991 |
| HA43 | 0.79639 |
| HA44 | 0.85492 |
| HA45 | 0.85848 |
| HA46 | 0.80687 |
| HA47 | 0.85238 |
| HA48 | 0.91196 |
| HA49 | 0.84027 |
| HA>49 | 0.7782 |
| **Average** | **0.82845** |

**Supplementary Figure 2 | Splitting the *refined training* set in 32 molecular size-related subsets**

Logistic regression submodels were trained on 32 subsets split by number of heavy atoms of compounds included in the *refined training* set. The curve of MCC obtained by 10-fold cross-validation is sufficiently flat and the average (MCC=0.83, values for each subset given on the right table) is very close to those of the *refined training* taken globally.

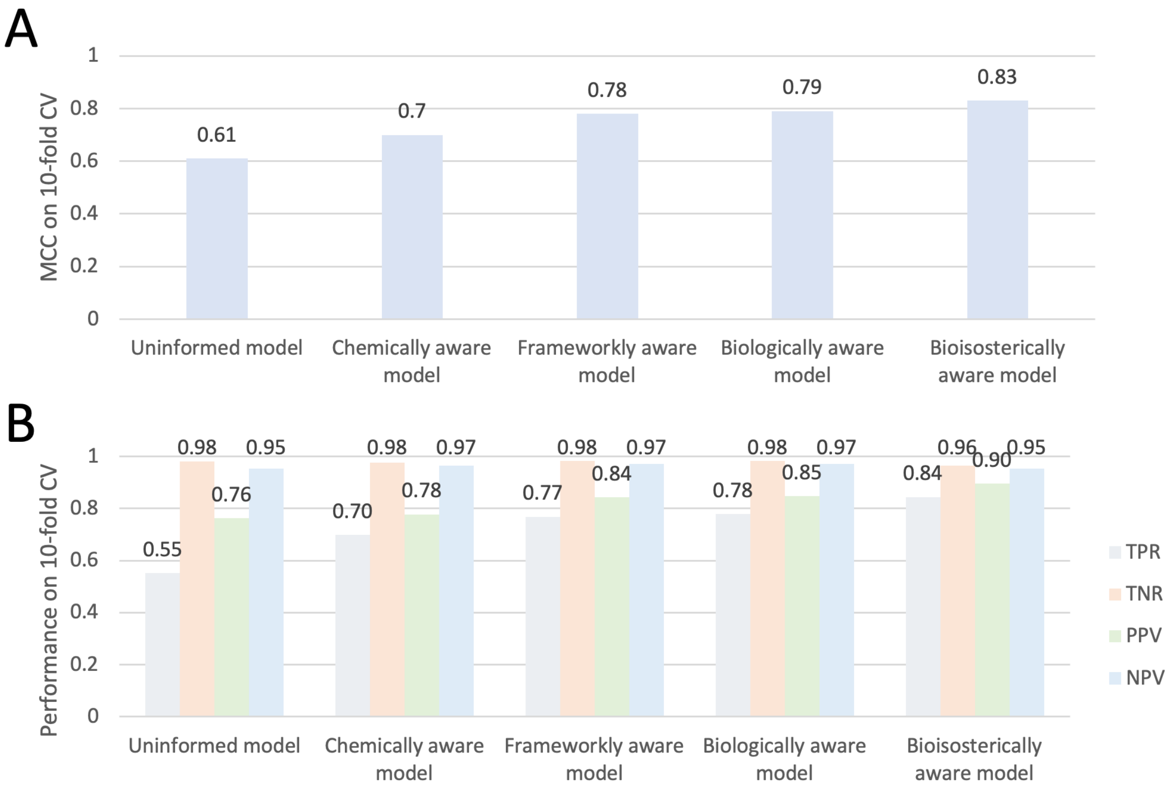

Supplementary Figure 3 | Robustness of the models built on the different training sets evaluated by 10-fold crossvalidation

[A] 10-fold crossvalidation Matthews Correlation Coefficient (MCC) of the logistic models built on the different training sets. [B] True Positive Rate (TPR), True Negative Rate (TNR), Positive Predictive Value (PPV) and Negative Predictive Value (NPV) on the same training sets.

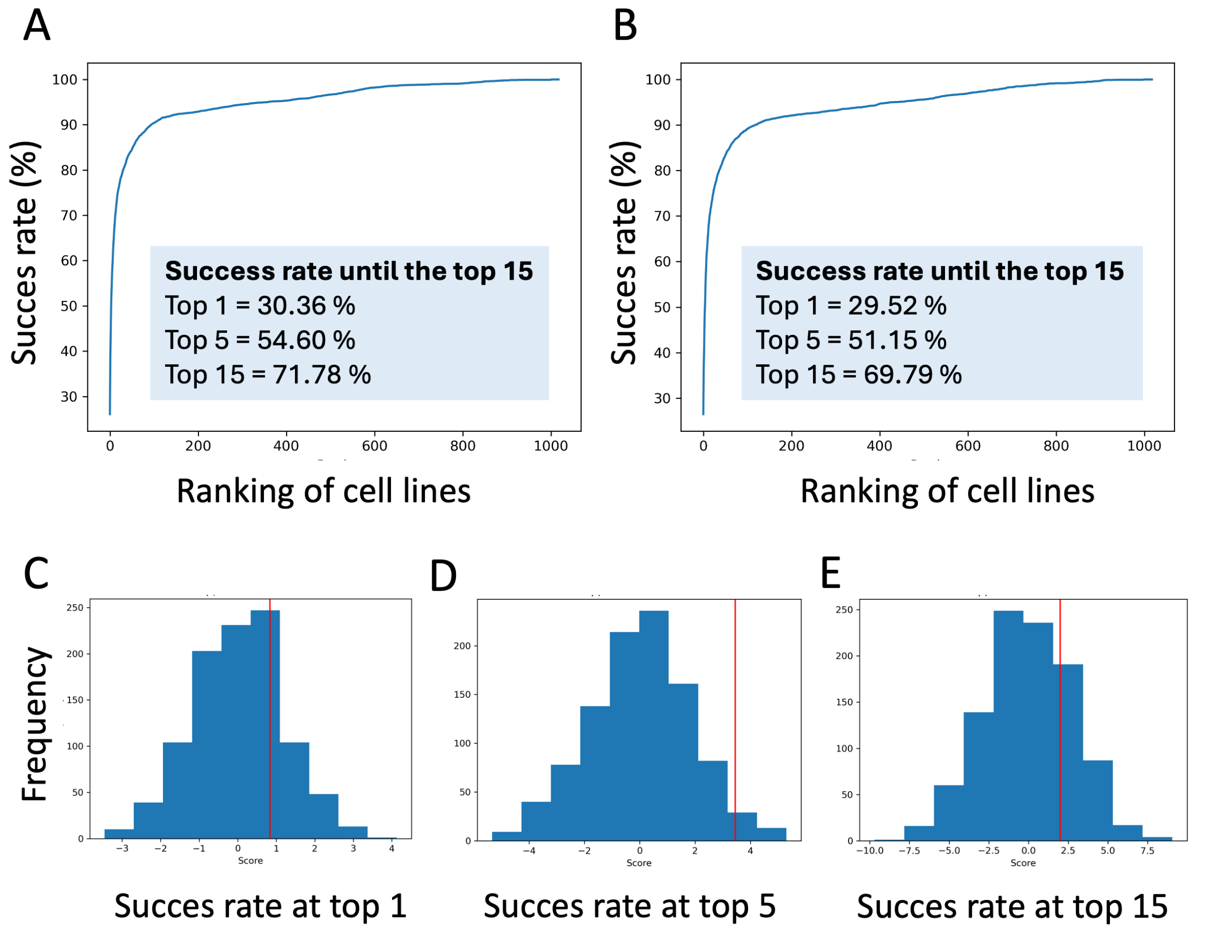

Supplementary Figure 4 | Predictive ability of reverse screening with respect to membrane permeation capacity prediction among.

Success rates in retrieving one experimentally validated cell line target on [A] 6516 test compounds predicted to have high permeation capacity. [B] 4465 compounds predicted to have low or moderate permeation capacity. [C-E] Each permutation test involves 1000 permutations of labels; P-values = 0.24 (top 1), 0.03 (top 5) and 0.24 (top 15).

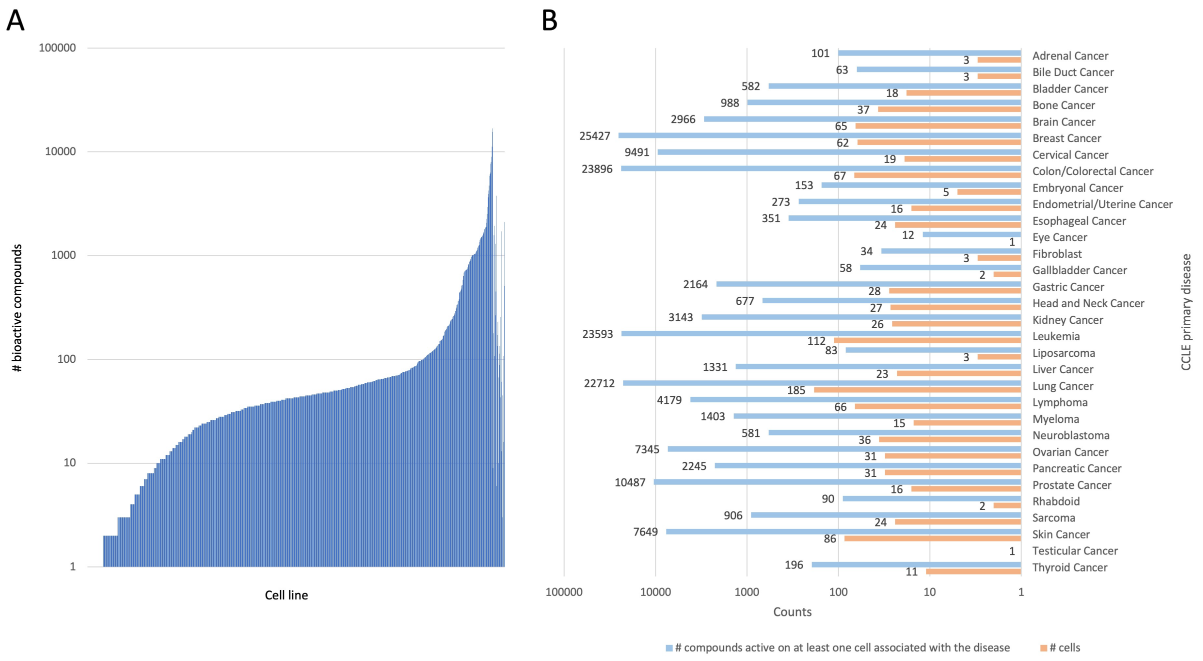

### Supplementary Figure 5 | Distribution of cytotoxic compounds by cancer cell line and disease

[A] Number of cytotoxic compounds by cancer cell line as reported in ChEMBL 29. [B] Number of cytotoxic compounds by annotated disease of cell lines (CCLE primary disease annotation). In blue the number of active compounds on at least one cell associated with the disease and in orange the number of cells.
